# Reduced miR-184-3p expression occurring in Type 2 diabetic pancreatic islets protects β-cells from lipotoxic and proinflammatory apoptosis via a CRTC1-dependent mechanism

**DOI:** 10.1101/2021.01.04.425234

**Authors:** Giuseppina E. Grieco, Noemi Brusco, Laura Nigi, Caterina Formichi, Daniela Fignani, Giada Licata, Lorella Marselli, Piero Marchetti, Laura Salvini, Laura Tinti, Agnese Po, Elisabetta Ferretti, Guido Sebastiani, Francesco Dotta

## Abstract

Loss of functional β-cell mass in Type 2 diabetes (T2D) involves molecular mechanisms including β-cell apoptosis, dysfunction, and/or dedifferentiation. MicroRNA miR-184-3p has been demonstrated to be involved in multiple β-cell functions including insulin secretion, proliferation and survival. However, downstream targets and upstream regulators of miR-184-3p have not yet been fully elucidated. Here, we showed that levels of miR-184-3p are reduced in human T2D pancreatic islets and that its reduction protected β-cells from lipotoxic- and inflammatory-induced apoptosis. Interestingly, CREB-Transcriptional Coactivator-1 (CRTC1) is a direct target of miR-184-3p and indeed its expression is upregulated in human T2D pancreatic islets. The downregulation of miR-184-3p in β-cells induced the upregulation of CRTC1 both at mRNA and protein level. Of note, miR-184-3p protection effect was dependent on CRTC1, since its silencing in human β-cells abrogates the protective mechanism exerted by miR-184-3p inhibition. Additionally, we found that the β-cell specific transcription factor NKX6.1, whose DNA binding sites were predicted to be present in human and mouse MIR184 gene promoter sequence, was reduced in T2D human pancreatic islets, in line with miR-184-3p downregulation, and was positively correlated with microRNA expression. Using chromatin immunoprecipitation analysis and mRNA silencing experiments, we demonstrated that NKX6.1 directly controls both human and murine miR-184 expression.

In conclusion, we found that miR-184-3p expression is controlled by the β-cell specific transcription factor NKX6.1 and that miR-184-3p reduction protects β-cells from apoptosis through the upregulation of its target gene CRTC1.

## Introduction

Type 2 diabetes mellitus (T2D) is a metabolic disease caused by the combinatorial effects of genetic background and environmental factors. The exposure to gluco-lipotoxic and inflammatory stresses progressively leads to β-cells failure, determining a significant reduction of functional β-cell mass and alteration of glucose homeostasis, thus resulting in chronic hyperglycaemia^1–3^.

Although β-cell mass has been shown to be significantly reduced in T2D patients due to apoptotic effects of gluco-lipotoxic stressors and of some inflammatory mediators^1–3^, some authors recently hypothesized and demonstrated an overestimation of such phenomenon^4^, indicating that additional not yet clarified mechanisms are involved in the pathogenesis of T2D^4^. Indeed, several studies reported that dedifferentiation of β-cells leading to the loss of β-cell phenotype characterizes T2D pathogenesis and actively contributes to β-cell functional deficit ^5^.

Interestingly, recent evidences suggested that dedifferentiation is accompanied by the acquisition of a protected-phenotype, rendering β-cell more resistant to metabolic and/or inflammatory stresses, though dysfunctional ^5,6^. These results are reinforced by the notion that β-cell failure, through dysfunction or dedifferentiation, can be apparently reversed ^7–9^, thus indicating that a subset of β-cells is dysfunctional but ready to potentially re-acquire their function^9–11^.

MicroRNAs (miRNAs), a class of small endogenous RNAs that negatively regulates gene expression^12,13^, have been reported to be pivotal modulators and rheostats of β-cell differentiation^14,15^, survival ^16^, phenotype maintenance ^17^ and function^18,19^.

Among these, miR-184-3p is one of the most enriched miRNA in pancreatic islets and in β cells^20^. Previous studies reported a reduced expression of miR-184-3p in pancreatic islets from both T2D animal models (db/db and ob/ob) ^21^ as well as in human pancreatic islets from T2D donors^20^. Several lines of evidence highlighted a potential role of miR-184-3p in β-cell protection mechanisms during metabolic insults typical of T2D^20^. Tattikota et al. demonstrated that miR-184-3p downregulation modulates Argonaute 2 (Ago2) expression, consequently leading to a compensatory β-cell expansion through indirect modulation of miR-375 activity^20^. Additionally, it has been reported that miR-184-3p downregulation is able to protect murine β-cell line MIN6 from metabolic and pro-inflammatory cellular stresses, depicting an antiapoptotic role for miR-184-3p ^21,22^. Furthermore, miR-184-3p regulates insulin secretion through direct targeting and expression regulation of solute carrier family 25 member 22 (Slc25A2), a mitochondrial glutamate carrier involved in glutamate transport across inner mitochondrial membrane ^23^.

Although a recent report ^24^ demonstrated that miR-184-3p is also regulated by the glucose sensor AMPK, the exact molecular mechanisms which confers protection to β-cells driven by miR-184-3p have not yet fully elucidated.

Here we provide further evidence that miR-184-3p is downregulated in human T2D pancreatic islets and that its reduction is due to the loss of the expression of β-cell phenotype characterizing transcription factor NKX6.1. Low levels of miR-184-3p confers protection to β-cells from metabolic/inflammatory stressful insults through the regulation of the transcription factor CREB Transcriptional Co-activator 1 (CRTC1).

## Materials and methods

### Human pancreatic islets

Human pancreatic islets were obtained from 13 non diabetic (gender: 7M, 6F; age 63.1±11.9y; BMI 27.1±3.8 Kg/m^2^) and 9 type 2 diabetic (T2D) multiorgan donors (gender: 6M; 3F; age 72.2±8y; BMI 26±1.9 Kg/m^2^) (see ***Supplementary Table 1***). Briefly, purified islets were prepared by intraductal collagenase solution injection and density gradient purification, as previously described^25^. At the end of the isolation procedure, fresh human pancreatic islets preparations were resuspended in CMRL culture medium (cat. 11-530-037, Fisher Scientific, Pittsburgh, PA, USA) supplemented with L-Glutamine 1% (cat. G7513-100ml), Antibiotic/Antimicotic 1% (A5955-100ML, Sigma Aldrich, St. Louis, MO, USA), FBS 10% and cultured at 28°C in a 5% CO_2_ incubator.

### Cell culture and transfection

#### EndoC-βH1 cells

EndoC-βH1 human β-cell line ^26,27^ was obtained from UniverCell-Biosolutions (Toulouse-France) and used for all the experiments between passages 68-80. EndoC-βH1 cells were cultured in coated flasks (coating medium: DMEM cat. 51441C, Penicillin/Streptomycin 1 % cat. P0781, ECM 1% cat. E1270 and Fibronectin from bovine plasma 0.2% cat. F1141-all from Sigma Aldrich, St. Louis, MO, USA) and cultured DMEM (cat. D6046) supplemented with 2% BSA fraction V (cat. 10775835001), β-Mercaptoethanol 50 μM (cat. M7522), L-Glutamine 1% (cat. G7513), Penicillin/Streptomycin 1% (cat. P0781), Nicotinamide 10 mM (cat. N0636), Transferrin 5.5 μg/mL (cat. T8158) and Sodium selenite 6.7 ng/mL (cat. S5261); all reagents are from Sigma Aldrich (St. Louis, MO, USA).

In order to evaluate CRTC1 mRNA and protein modulation by hsa-miR-184-3p, EndoC-βH1 cells were plated at a density of 1.37L10^5^ cells/cm^2^ in 24 or 6 well plates. After 24h, EndoC-βH1 cells were transfected with 100 nM of miRVANA miR-184 inhibitor (cat. 4464084, ID: MH10207-Invitrogen, Waltham, MA, USA) or with 100 nM negative control inhibitor (cat. 4464076-Invitrogen, Waltham, MA, USA) using Lipofectamine 2000 (cat. 11668-019, Invitrogen, Waltham, MA, USA).

NKX6.1 was silenced using specific siRNA as follows. EndoC-βH1 cells were plated at a density of 1.37□10^5^ cells/cm^2^ in 24 or 6-well plates and were transfected after 24h by using Lipofectamine 2000 (cat. 11668-019, Invitrogen, Waltham, MA, USA) for 48h with 25 nM of CTR siRNA (cat. D-001810-10-05 ON-TARGET plus non-targeting pool siRNA 5 nmol, Dharmacon- Lafayette, CO, USA) or with 25 nM of NKX6.1 siRNA (cat. L-HUMAN-XX-0005, ID. L-020083-00-0005, ON-TARGET plus SMART pool 5 nmol, Dharmacon- Lafayette, CO, USA).

EndoC-βH1 cells were subjected to lipotoxic or inflammatory stimuli as previously described ^28–30^. Lipotoxic stress have been induced by 2 mM of sodium palmitate (cat. P9767-5G, Sigma Aldrich, St. Louis, MO, USA) for 24h or with EtOH 0,5% as control treatment. Pro-inflammatory stress was induced using a cytokines mix composed of: IL-1β (50 U/mL) (cat. 201-LB-005, R&D system Minneapolis, MN, USA), TNFα (1000 U/mL) (T7539 from Sigma Aldrich, St. Louis, MO, USA) and IFNγ (1000 U/mL) (cat. 11040596001 from Roche, Basel, Switzerland) for 24h.

In order to evaluate the role of CRTC1 and hsa-miR-184-3p in β-cells protection and survival during lipotoxic or pro-inflammatory stresses, EndoC-βH1 cells were plated at a density of 1.37□10^5^ cells/cm^2^ in 24 or 6 well plates and transfected after 24h using Lipofectamine 2000 (cat. 11668-019, Invitrogen, Pittsburgh, PA, USA) with different combinations of CRTC1 siRNA and/or miR-184 inhibitor: 100 nM negative control miRNA inhibitor (cat. 4464076-Invitrogen, Pittsburgh, US) + 50 nM of CTR siRNA (cat. D-001810-10-05 ON-TARGET plus Non-targeting Pool siRNA 5 nmol, Dharmacon-Lafayette, CO, USA); 100 nM negative control miRNA inhibitor + 50 nM of CRTC1 siRNA (cat. L-HUMAN-XX-0005, ID L-014026-01-0005, ON-TARGET plus SMART pool 5 nmol, Dharmacon-Lafayette, CO, USA); 100 nM of miRVANA miR-184 inhibitor (cat. 4464084, ID. MH10207-Invitrogen, Pittsburgh, MO, USA) + 50 nM of CTR siRNA; 100 nM of miRVANA miR-184 inhibitor (cat. 4464084, ID. MH10207-Invitrogen, Pittsburgh, US) + 50 nM of CRTC1 siRNA. After 24h, transfected EndoC-βH1 cells were subjected to lipotoxic or pro-inflammatory stresses as reported above, and then analyzed to evaluate pyknotic nuclei, active Caspase-3 and CRTC1 mRNA and protein expression.

#### MIN6 cells

MIN6 cells were obtained from AddexBio (San Diego, CA, USA), cultured between passages 11-25 and maintained as previously described^31^. MIN6 were cultured in DMEM (cat. 5671, Sigma-Aldrich, St. Louis, MO, USA) supplemented with L-Glutamine 1%, Antibiotic/Antimicotic 1% (A5955, Sigma Aldrich, St. Louis, MO, USA), FBS 15% (cat. ECS0180L, Euroclone, Milan, Italy), sodium pyruvate 1 % (cat. ECM0542D, Euroclone, Milan, Italy) and β-Mercaptoethanol 50 μM. Lipofectamine 2000 has been used to transfect MIN6 cells with CRTC1 overexpressing plasmid (EX-Mm18750-Lv183) or with control plasmid (EX-EGFP-Lv151) (both from Genecopoeia, Rockville, MD, USA). Lipotoxic stress has been induced by using 0.5 mM of sodium palmitate (Sigma-Aldrich, St. Louis, MO, USA) for 48h as previously described^32^. Pro-inflammatory stress was induced by a cytokine mix composed of IL-1β (5 ng/mL, cat. I5271-Sigma Aldrich, St. Louis, MO, USA), TNFα (30 ng/mL, cat. T7539) and IFNγ (10 ng/mL, cat. 485MI- R&D System, Minneapolis, MN, USA) for 24 h. Oxidative stress was induced using H_2_O_2_ (cat. H1009, Sigma Aldrich, St. Louis, MO, USA) 100 μM for 90min as previously described.

#### 1.1B4 cells

1.1B4 human pancreatic β-cell line^33^ (obtained from Sigma-Aldrich) was cultured in RPMI (cat. R0883, Sigma Aldrich, St. Louis, MO USA) supplemented with L-Glutamine 1%, Antibiotic/Antimycotic 1%, FBS 10%. Transfection was performed by using Lipofectamine 2000 with 100 nM miRVANA miR-184 inhibitor (cat. 4464084, ID: MH10207-Invitrogen, Pittsburgh, PA, USA) respect to 100 nM negative miRNA control inhibitor (cat. 4464076-Invitrogen, Pittsburgh, PA, USA).

#### HeLa cells

HeLa cells were cultured in DMEM (cat. 5671, Sigma-Aldrich, St. Louis, MO, USA) supplemented with L-Glutamine 1%, Antibiotic/Antimycotic 1%, FBS 10%. Lipofectamine LTX (cat. 15338100, Thermo Fisher Scientific, Waltham, MA, USA) has been used for HeLa cell line transfection with plasmid DNA or with 100nM of miRVANA miR-184 inhibitor or 100 nM negative control miRNA inhibitor (cat. 4464076-Invitrogen, Waltham, MA, USA).

### Predictive analysis of hsa-miR-184-3p target genes

TargetScan Human 7.0 (www.targetscan.org) algorithm was used to retrieve the list of conserved hsa-miR-184-3p predicted target genes (***Supplementary Table 2***).

### Dual luciferase reporter assay

The luciferase reporter vector HmiT060400a-MT06, containing wild type (WT) CRTC1 3′UTR sequence, and the vector HmiR0259-MR04, containing hsa-miR-184-3p precursor, were obtained from Genecopoeia (Rockville, MD, USA). HeLa cells were plated at a density of 4×10^4^/well in 24-well plates and, after 24h, transiently co-transfected with 245□ng of hsa-miR-184-3p precursor plasmid vector and 5□ng of CRTC1 luciferase reporter vector, using 1,75μl Lipofectamine LTX transfection reagent, for 48 h. After transfection, luciferase activity was measured at luminometer Glomax 20/20 using Dual-Luciferase Reporter Assay System (cat. #E1910-Promega, Fitchburg, USA) following manufacturer's instructions. Data of Firefly luciferase activity in frame with CRTC1-3’UTR sequence were normalized to Renilla luciferase activity. Specific experimental points were used to extract total RNA using miRNeasy mini kit (cat. 217004-Qiagen, Hilden, Germany) in order to evaluate miR-184-3p upregulation upon transfection.

### MiRNAs and genes qRT-PCR analysis

For miRNA and gene qRT-PCR expression analysis, total RNA including small RNAs <200nt was extracted with miRNeasy mini kit (cat. 217004-Qiagen, Fitchburg, Germany). Total RNA was quantified using Qubit 3000 Fluorometer (Thermo Fisher Scientific, Waltham, MA, USA) and quality evaluated using 2100 Bioanalyzer-RNA 3000 Pico Kit (Agilent Technologies), excluding those samples with RIN<5.0. For miRNA expression analysis, cDNA was prepared using TaqMan™ MiRNA Reverse Transcription Kit (cat. 4366596-Invitrogen, Waltham, MA, USA), while cDNA from mRNAs were synthesized using SuperScript™ III First-Strand Synthesis System (cat. 18080044-Invitrogen, Waltham, MA, USA) alongside with random hexamers. Real-Time PCR analysis was performed using Taqman miRNA expression assays or Taqman gene expression assays using the primers listed in ***Supplementary Table 3***, following manufacturer’s recommendations. Data were collected and analyzed through Expression Suite software 1.0.1 (Thermo Fisher Scientific, Waltham, MA, USA) using 2^-ΔCt^ or 2^-ΔΔCt^ method. VIIA7 Real-time PCR instrument was used to analyze both miRNA and gene expression.

### Gene Expression Omnibus (GEO) dataset analysis

Gene Expression Omnibus (GEO) database (https://www.ncbi.nlm.nih.gov/gds) was interrogated to evaluate the expression of miR-184-3p predicted target genes in T2D pancreatic islets. We looked for previously published studies reporting genes expression profiling performed on pancreatic islets from T2D donors. We retrieved data from GSE20966 and GSE25724 datasets ^34,35^ reporting the analyzed and normalized results obtained from genes microarray profiling of pancreatic islets or LCM isolated β-cells from non-diabetic and T2D donors. Specifically, expression data corresponding to miR-184-3p predicted target genes were obtained and plotted by using Graphpad 8.0 software. Mann-Whitney U test was used to analyze statistical significance (p<0.05).

### Western blot analysis

Total protein from EndoC-βH1, MIN6, 1.1B4 or HeLa cells were extracted using RIPA lysis buffer (50mM Tris pH 7.6, 0.5% Sodium Deoxycholate, 1% IGEPAL, 0.1% SDS, 140mM NaCl, 5mM EDTA) supplemented with 1X protease inhibitors (Roche, Basel, Switzerland). Total proteins were quantified using Bradford assay and 50μg protein/lane were separated using SDS-PAGE Tris-Glycine gradient Bis-Acrylamide gel 4-20%. Proteins were then transferred to PVDF membrane using wet electrophoresis system. Upon transfer, PVDF membranes were washed 3 times with PBST1X and then incubated 2h with 5% non-fat dry milk in PBST1X. To identify CRTC1, rabbit monoclonal anti-CRTC1 (cat. ab92477, Abcam) was diluted 1:1000 in 5% non-fat dry milk in PBST1X and incubated o/n at 4°C and then with Goat anti-rabbit (cat. SC-2004 – Santa Cruz, Dallas, TX, USA) diluted 1:5000 in 2% non-fat dry milk in PBST1X 1h RT. NKX6.1 was detected using rabbit monoclonal anti-NKX6.1 (cat. 54551, Cell Signaling Technology, Danvers, MA, USA) diluted 1:500 in 5% non-fat dry milk in PBST1Xand incubated o/n at +4°C and then with Goat anti-rabbit-HRP (Santacruz, Dallas, TX, USA- SC-2004) diluted 1:5000 in 2% non-fat dry milk in PBST1X 1h RT. All blots were normalized using β-actin; mouse monoclonal anti-β-actin (cat. A5441, Sigma Aldrich, St. Louis, MO, USA) was diluted 1:10000 in 5% non-fat dry milk in PBST1X and incubated 1h at RT and subsequently with Goat anti-mouse-HRP (Santa Cruz, Dallas, TX, USA-A1303) diluted 1:1000 in 2% non-fat dry milk in PBST1X 30min RT.

Chemiluminescent signal was detected by using ECL solution (GE Healthcare, Little Chalfont, Buckinghamshire, UK). Chemiluminescent analysis of immunoblot results was performed by densitometric analysis using LAS400 analyzer (GE Healthcare, Little Chalfont, Buckinghamshire, UK-RPN2232) and Image J software.

### CRTC1 targeted Mass Spectrometric (MS) analysis

To perform CRTC1 targeted MS analysis, EndoC-βH1 cells were prepared as previously described^36,37^. Briefly, cells were lysed with RIPA buffer and protein lysate was quantified through BCA assay. Then, protein lysate was mixed with 400 μL of urea 8M in Tris-HCl 100nM pH 8,5 (UA), with the addition of 100 mM DTT. The mixture was loaded on a filter 10 K Pall, incubated 30min RT and centrifuged at 13,800xg for 30min. Filter was then washed twice with 400 μL of UA and centrifuged at 13,800xg for 30min, then incubated with 100 μL of 50 mM of iodoacetamide (IAC) solution in a thermo-mixer for 1min 150 rpm and without mixing for 20min, then centrifuged at 13,800xg for 20min. Filter was then washed twice with 400 μL of UA and centrifuged at 13,800xg for 30min, twice with 400 μL of 50 mM ammonium bicarbonate (AMBIC) and centrifuged twice at 13,800xg for 30min and for 20 min, respectively. Then 40 μL of 50 mM AMBIC were added to the filter together with trypsin (ratio trypsin/proteins 1:25) and incubated O/N at 37°C. Then, the sample was transferred in a collecting tube and centrifuged at 13,800xg for 10min. Subsequently, 100 μL of 0.1% formic acid were added on the filter and centrifuged 13,800xg 10min. Finally, filter was discarded and solution was desalted with OASIS cartridges according to manufacturer’s instructions. Then, peptides were concentrated using SpeedVac and sample was resuspended in a solution of 3% acetonitrile, 96.9% H_2_O and 0.1% formic acid. Analyses were performed on a Q-Exactive Plus mass spectrometer (Thermo Fisher Scientific, Waltham, MA, USA), equipped with electrospray (ESI) ion source operating in positive ion mode. The instrument is coupled to an UHPLC Ultimate 3000 (Thermo Fisher Scientific, Waltham, MA, USA). The chromatographic analysis was performed on a column Acquity UPLC Waters CSH C18 130Å (1 mm X 100 mm, 1,7 μm, Waters) using a linear gradient with 0.1% formic acid in water (phase A) and 0.1% formic acid in acetonitrile (phase B). The flow rate was maintained at 100μl/min and column oven temperature at 50°C. Mass spectra were recorded in the mass to charge (m/z) range 200-2000 at a resolution of 35K at m/z 200. The mass spectra were acquired using a “data dependent scan”, able to acquire both the full mass spectra in high resolution and to “isolate and fragment” the ten ions with highest intensity present in the full mass spectrum. The raw data obtained were analyzed using the Biopharma Finder 2.1 software from ThermoFisher Scientific. The elaboration process consisted in the comparison between the peak list obtained “in silico” considering the expected aminoacid sequence of human CRTC1 protein (Uniprot ID: Q6UUV9), trypsin as digestion enzyme and eventual modifications (carbamidomethylation, oxidation, etc.). Briefly, for quantitative analysis, peptides with the highest reliability were manually chosen for CRTC1, β-ACT and CHGA proteins. Then, aminoacid sequences of peptides of interest were identified in the raw spectrum to collect m/z and NL values. To obtain fold change, NL values of CRTC1 were normalized for NL values of both β-ACT and CHGA. One-Way ANOVA multiple comparison for statistical analysis and significance cutoff was set as p value<0,05.

### Apoptosis detection

Apoptosis rate was detected by cytofluorimetric analysis of cleaved Caspase-3^+^ cells and by pyknotic nuclei count. To detect the levels of cleaved Caspase-3, cells were fixed and permeabilized with BD Perm/Wash (cat. 554723-BD Biosciences, Franklin Lakes, New Jersey) and Cytofix/Cytoperm (cat. 554722-BD Biosciences, Franklin Lakes, New Jersey) buffers and stained by using 10 μg/ml of rabbit polyclonal anti-cleaved Caspase-3 (cat. ab13847, Abcam) antibody [or 10 μg/ml of rabbit IgG Isotype Control (cat. ab199376 Abcam)] and 1:500 diluted fluorescent secondary antibody Goat anti-rabbit Alexa Fluor 594 (cat. A11037, Thermo Fisher Scientific, Waltham, MA, USA). Cells were analyzed in a BD FACS Canto flow cytometer with FACS Diva software (Franklin Lakes, New Jersey). Then, FlowJo software was used for further analysis of Caspase-3^+^ cells. Pyknotic nuclei count was performed using Hoechst 33342 (cat. 62249-Invitrogen, Waltham. MA, USA) diluted 1:100 and added 1 μl/well in 24 well plates; manual counting was performed to detect the total number of pyknotic nuclei was normalized for number of total nuclei.

### Predictive analysis of miR-184 promoter regulation

MatInspector algorithm has been used to identify the transcription factors putatively having one or more recognition sites on DNA sequence corresponding to the miR-184 proximal promoter, 500 bp upstream of transcriptional starting site (TSS) of miR-184. The sequence 500 bp upstream of TSS of miR-184, corresponding to its proximal promoter as already described ^38^, was retrieved through Ensembl database for human (ENST00000384962) and murine sequence (ENSMUST00000083662), and selected to perform bioinformatic analysis.

### Immunofluorescence

Cultured MIN6 cells were immunostained as follows: treated or not-treated cells were fixed in 4% PFA for 10 min, permeabilized in 0,25% Triton-X-100 for 5 min and blocked in 3% BSA+0.05% Triton-X_100 in PBS for 30 min. MIN6 cells were incubated with primary antibody rabbit monoclonal anti-NKX6.1 (cat. 54551, Cell Signaling Technology, Danvers, MA, USA) diluted 1:100 in BSA 1% in PBS for 1h, rinsed with PBS and incubated with goat anti-rabbit-488 1:500 in 1% BSA in PBS 1:500 was used as secondary antibody.

### Chromatin Immunoprecipitation (ChIP) analysis

ChIP analysis was performed using MagnifyTM Chromatin Immunoprecipitation System (cat. 492024-Thermo Fisher Scientific, Waltham, MA, USA) following manufacturer’s instructions. Briefly, cultured MIN6 and EndoC-βH1 cell lines were incubated with formaldehyde 1% for 10min. Fixation was stopped at room temperature with 114 μl of 1,25 M Glycine and then samples were washed with PBS and 199 μl of Lysis Buffer in the presence of 1 μl of Protease Inhibitor 200X, at 4°C. Chromatin shearing was performed through sonication. The sonicated fragments were verified on a 2% TAE agarose gel. Sonicated chromatin was diluted and added to 0.2 ml PCR tube containing primary antibodies (3ug of rabbit anti-NKX6.1 cat. 54551 (Cell Signaling Technology, Danvers, MA, USA); 3 μg of rabbit-anti-acetyl-histone H3-cat. 06599, Millipore, St. Louis, MO, USA), coupled to the protein A/G Dynabeads. Rabbit IgG isotype control antibody (cat. 02-6102, Invitrogen, Waltham, MA, USA) was used as negative control. Samples were incubated 2h at 4°C. After incubation, the chromatin was washed with IP Buffer 1 and IP Buffer 2 provided by kit. Following the last wash, Reverse Crosslinking Buffer + Proteinase K was added to all samples, followed by incubation at 55°C for 15 minutes; specific antibodies-immunoprecipitated and not immunoprecipitated chromatin (referred as input control) was then recovered and incubated at 65°C for 15 minutes. Samples were incubated with DNA Purification Buffer + DNA Purification Magnetic Beads and then washed with DNA Wash Buffer and DNA Elution Buffer. After incubation at 55°C for 20 minutes, samples were stored at −20°C.

### Real-Time PCR for ChIP analysis of miR-184 promoter

Real-Time PCR was performed to quantify the amplification of immunoprecipitated miR-184-3p sequence promoter fragment. 1/30 recovered DNA was loaded into the PCR reaction composed of: 12.5 μl Maxima SYBR Green qPCR Master mix (2X), 0.05 μl ROX solution and nuclease-free H2O. The following primers were used for the miR-184 human promoter (300 nM each): forward primer 5’-AATGGCATGTGGGTGTTGGT-3’; reverse primer 5’-AGGGCTCCTGCAGGTCTGA-3’. The following primers were used for miR-184-3p murine promoter (500 nM): forward primer 5’AATGGCATGTGGGTGTTGGT 3’; reverse primer 5’ AGGGCTCCTGCAGGTCTGA 3’.

The reaction was incubated at 95°C for 10min, 95°C for 15 sec – 60°C for 30sec-72°C for 30sec (40 cycles) in ViiA7 Real Time PCR instrument (Thermo Fisher Scientific, Waltham, MA, USA). Data were collected and analyzed through Expression Suite software 1.0.1 (Thermo Fisher Scientific, Waltham, MA, USA) using 2^-ΔCt^ method. Raw Ct data resulting from specific antibodies-immunoprecipitated chromatin were normalized to negative isotypic control rabbit IgG (cat. 492024-Thermo Fisher Scientific, Waltham, MA, USA).

## Results

### miR-184-3p is downregulated in human Type 2 Diabetic (T2D) pancreatic islets and protects β-cells from apoptosis

MiRNA miR-184-3p has been demonstrated to play pivotal roles in fine tuning and modulation of islet/β-cell function. With the aim to further analyze miR-184-3p function and to confirm previous findings, we firstly analyzed miR-184-3p expression in human pancreatic islets isolated from pancreata derived from n=13 non-diabetic and n=9 T2D multiorgan donors (**Supplementary Table 1**). We detected a significant reduction of miR-184-3p expression in pancreatic islets isolated from T2D vs non-diabetic donors (−2.9 fold change vs CTR, p=0.043 non-parametric Mann-Whitney U test) (**Fig. 1a**). No correlation of donors age or BMI with miR-184-3p expression levels was detected (data not shown).

**Figure 1.**
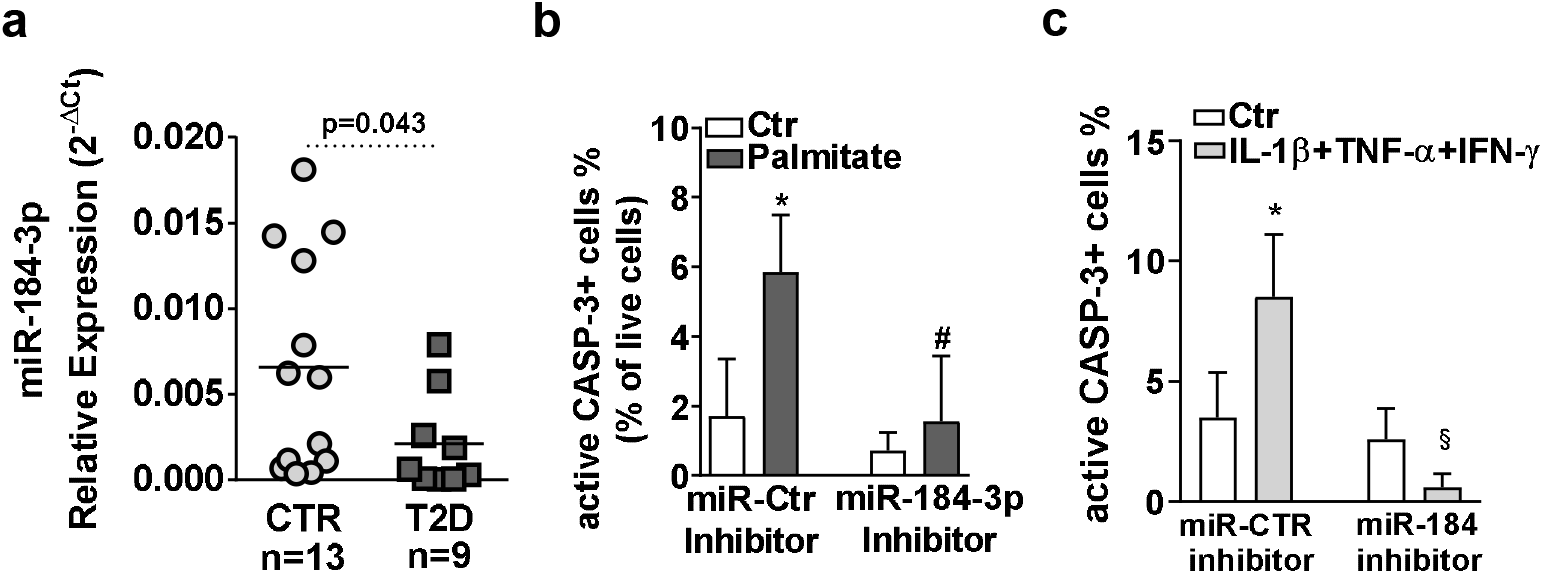
miR-184-3p is downregulated in T2D human pancreatic islets and protects β-cells from palmitate- and cytokines-induced apoptosis. **(a)** RT-Real-Time PCR analysis of miR-184-3p expression in T2D human pancreatic islets (n=9; dark grey squares) and in non-diabetic (n=13; light grey circles) multiorgan donors; data are shown as 2^-ΔCt^ relative expression single dot values normalized using RNU44 and RNU48, alongside with mean value; statistics using Mann-Whitney U Test. (**b)** Cytofluorimetric evaluation of active Caspase-3 positive (active-CASP-3+) EndoC-βH1 cells respectively after 48h palmitate treatment (* p=0.0008 vs. not treated miR-Ctr inhibitor; # p=0.0005 vs. palmitate treated miR-Ctr inhibitor transfected) and **(c)** after 24h cytokines mix (IL-1β+TNF-α+IFN-γ) treatment (* p=0.032 vs. not treated miR-Ctr inhibitor; § p<0,0001 vs cytokines treated miR-Ctr inhibitor). Data are reported as mean ± SD of percentage (%) of positive cells on total live cells. Statistics performed using ANOVA analysis with Bonferroni’s multiple comparison test (n=4 independent experiments).

In pancreatic islets derived from T2D mouse models (*db/db*, *ob/ob*) and in murine β-cell lines (MIN6), miR-184-3p expression reduction has been proven to be involved in a pro-survival process, adopting a not elucidated protection mechanism^21^. To verify whether such process was occurring also in human β-cells, we inhibited miR-184-3p expression in the human β cell line EndoC-βH1 ^26^ exposed to palmitate or to cytokines mix (IL-1β+TNF-α+IFN-γ) to simulate lipotoxic or inflammatory stress. MiR-184-3p inhibition protected human β-cell from apoptosis upon 48h of palmitate exposure, as shown by active Caspase-3 staining (**Fig. 1b and Suppl. Fig.1a**) and confirmed by pyknotic nuclei count (**Suppl. Fig.1b**). Such protection was observed also upon exposure of EndoC-βH1 to inflammatory stimuli, using a mix of pro-inflammatory cytokines (IL-1β+TNF-α+IFN-γ) (**Fig. 1c and Suppl. Fig.1c-d)** demonstrating that miR-184-3p inhibition protect β-cells from both metabolic and inflammatory stress conditions.

Collectively, these data confirm that miR-184-3p expression is reduced in pancreatic islets derived from T2D multiorgan donors and that this molecular feature is involved in β-cells protection from palmitate- and cytokines-induced apoptosis.

### The predicted target of miR-184-3p, Creb Transcriptional Coactivator-1 (CRTC1), is upregulated in human LCM captured β-cells and in pancreatic islets from type 2 diabetic donors

In order to further elucidate the function of miR-184-3p, we reviewed its predicted target genes by using the algorithm TargetScan 7.1 (http://www.targetscan.org/vert_71/). We identified 29 high-ranked and conserved predicted hsa-miR-184-3p target genes (listed in **Supplementary Table 2)**. Since miR-184-3p downregulation should result into an increased expression of its target genes in islet/β-cell context, we interrogated a previously published and online available dataset reporting high-throughput gene expression data of laser capture microdissected non-diabetic and T2D human β-cells (GEO dataset: GDS3782-GSE20966)^34^, for the expression of miR-184-3p predicted target genes. Overall, 25/29 genes were detected in the samples analyzed in the dataset and 3/25 resulted significantly differentially expressed (**Fig. 2a and Supplementary file 1**). Among them, Creb Transcriptional Coactivator-1 (CRTC1) resulted upregulated in T2D vs non-diabetic β-cells (**Fig. 2b**) (CRTC1, p=0.0039-Mann Whitney U test) and, therefore, in line with reduced miR-184-3p expression in T2D pancreatic islets. A similar differential expression (p=0.02-Mann Whitney U test) was observed in another dataset (GEO dataset: GDS3882-GSE25724) which includes genes expression data from non-diabetic and T2D collagenase-isolated human pancreatic islets (**Fig. 2c**). Finally, we evaluated CRTC1 mRNA expression using RT-Real-Time PCR in a subset of isolated islets samples from our cohort of non-diabetic and T2D multiorgan donors (**Supplementary Table 1**). The analysis showed a similar trend in terms of differential expression (although not significant, p=0.07-Mann Whitney U test) observed in previous datasets (**Fig. 2d**).

**Figure 2.**
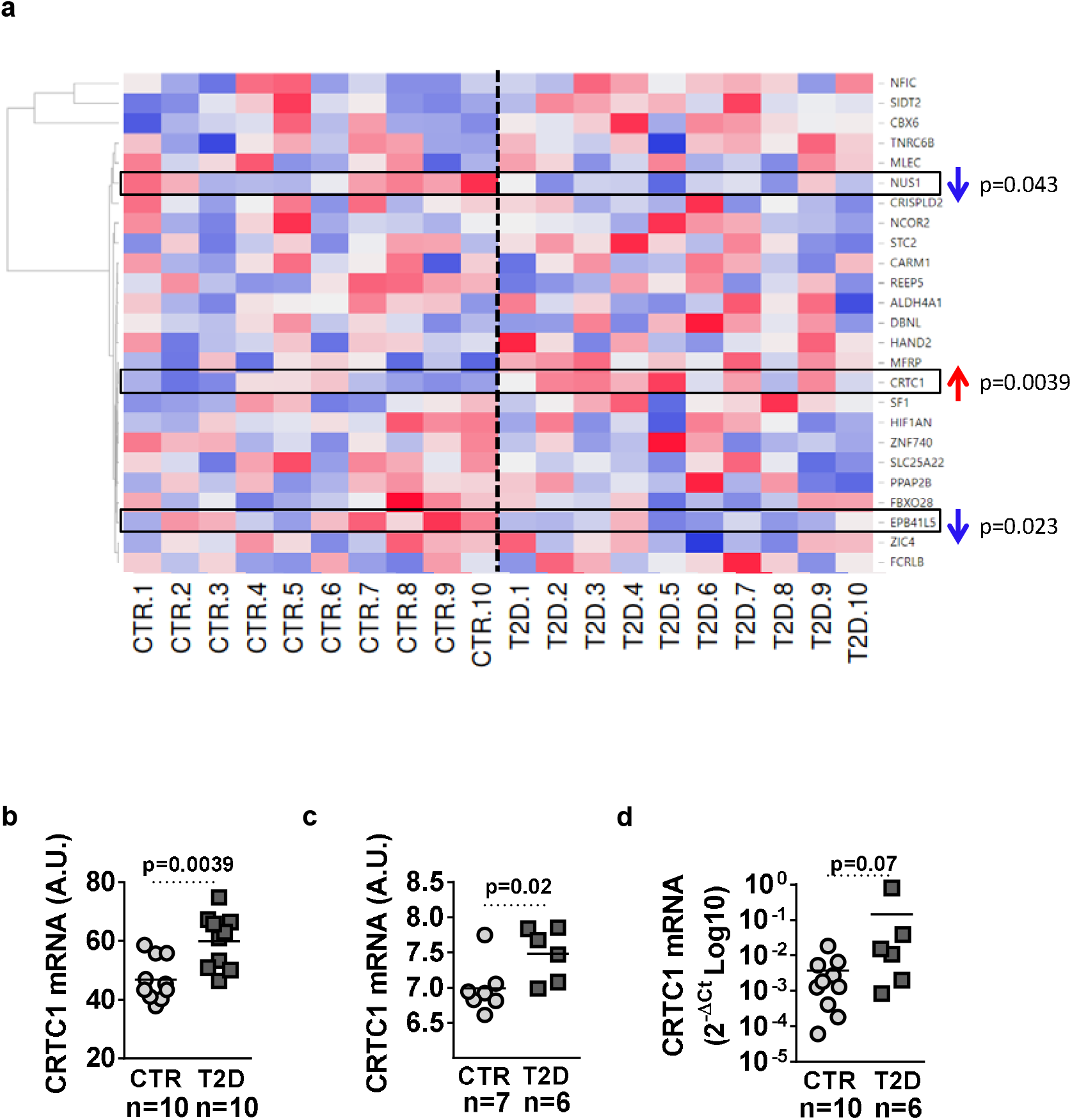
CRTC1 is upregulated in LCM captured human β-cells and in pancreatic islets from T2D donors. (**a)** Hierarchical clustering heatmap expression analysis of hsa-miR-184-3p predicted target genes (n=25) detected in LCM β-cells captured from n=10 T2D vs n=10 non-diabetic donors (data previously published and deposited in GEO dataset GDS3782-GSE20966). Gene expression values are reported as A.U. (Arbitrary Unit count) scale colors (high expression: red; low expression: blue). (**b**) Dot plot graph showing CRTC1 expression values analyzed in the dataset reported in (a). Values are reported as Arbitrary Units (A.U.) alongside with mean value. (**c)** Dot plot graph showing CRTC1 expression in isolated human pancreatic islets from n=6 T2D vs n=7 non-diabetic donors (data previously published and reported in GEO dataset GDS3882-GSE25724). Values are reported as A.U. alongside with mean value. (**d)** RT-Real Time PCR validation of CRTC1 expression in human pancreatic islets isolated from n=6 T2D vs n=10 non-diabetic multiorgan donors; data are reported as 2^-ΔCt^ relative expression single dot values normalized using β-ACT and GAPDH; statistics performed using Mann Whitney U test (p<0,05).

Such data uncover the potential presence of a miR-184-3p-CRTC1 regulatory axis in β-cell function and survival. Consequently, taking into account that previous reports attributed a critical role to CRTCs (Creb Regulated Transcriptional Coactivators) in the regulation of cell metabolism and survival through multiple signaling pathways ^39,40^, we aimed at establishing whether CRTC1 is a miR-184-3p target gene.

### MiR-184-3p directly modulates CRTC1 expression

Human CRTC1-3’UTR sequence showed 3 binding sites for miR-184-3p (**Fig. 3a**). Although these binding sites are widely conserved among primates, no binding site for miR-184-3p is present within the 3’UTR of murine CRTC1, despite the large homology observed between human and mouse CRTC1 coding sequence (>85%). Firstly, in order to assess the binding of miR-184-3p to 3’UTR of human CRTC1, we co-transfected HeLa cell line with a plasmid containing the entire 3’UTR sequence of CRTC1 in a Firefly-Renilla Luciferase vector, alongside with miR-184-3p expressing plasmid. Luciferase assay revealed that miR-184-3p overexpression induced a significant reduction of Firefly luciferase activity, thus confirming the interaction of miR-184-3p with CRTC1 3’UTR sequence (**Fig. 3b**). Subsequently, to assess miR-184-3p mediated CRTC1 expression modulation at mRNA and protein levels, we inhibited miR-184-3p for 24h or 48h in EndoC-βH1 and in 1.1B4 human β-cell lines as well as in HeLa cells, and then measured CRTC1 expression by RT-Real-time PCR and by Western Blot analysis. The inhibition of miR-184-3p led to a significant increase of CRTC1 mRNA expression at 24h and 48h post-transfection and a corresponding protein increase at 48h both in EndoC-βH1 (**Fig. 3c and Fig. 3d**) and in 1.1B4 β-cell line (**Suppl. Fig. 2a and Suppl. Fig. 2b**), while in HeLa cells, CRTC1 protein increase was observed as early as 24h post-transfection (**Suppl. Fig. 2c and Suppl. Fig. 2d**).

**Figure 3.**
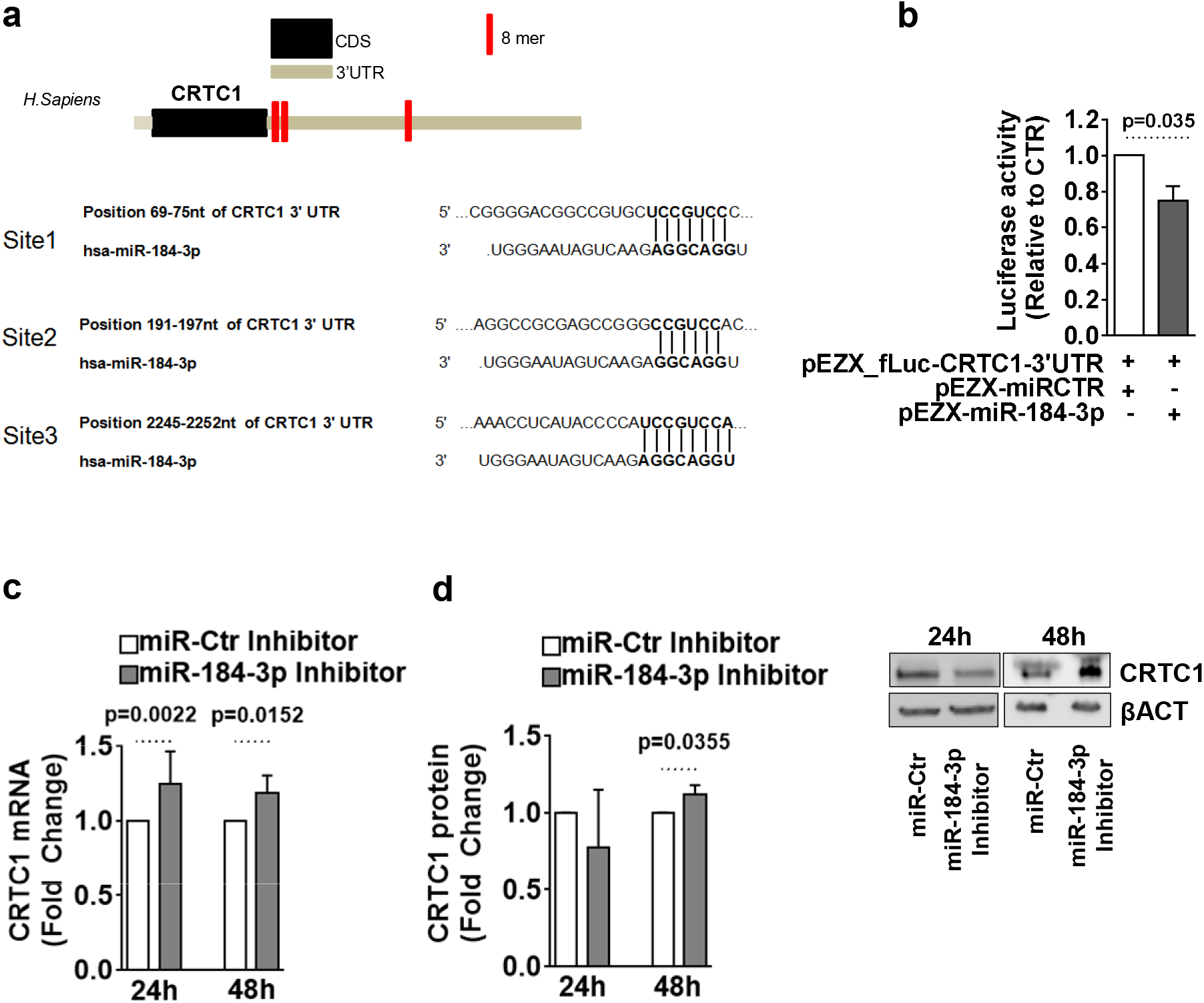
miR-184-3p directly modulates the expression of CRTC1. (**a,** *upper panel*) Graphical scheme depicting CRTC1 mRNA sequence (ENST00000338797.6) reporting 3’UTR and hsa-miR-184-3p predicted binding sites (in red); (**a, lower panel**) hsa-miR-184-3p binding sites nucleotide position and sequence context within 3’UTR region of CRTC1 mRNA. **(b)** Luciferase assay performed in HeLa cells transfected with hsa-miR-184-3p overexpressing plasmid respect to scrambled control vector transfected cells; values are shown as mean ± SD of relative luciferase activity respect to control, upon normalization of firefly luciferase unit to Renilla Luciferase activity; statistics using Mann Whitney U test (n=4). **(c)** RT-Real Time PCR analysis of CRTC1 expression in EndoC-βH1 cells transfected with a synthetic inhibitor of hsa-miR-184-3p; data are reported as fold change vs scrambled control; statistics using Mann Whitney U Test (n=6); (**d**) Western Blot analysis of CRTC1 (78 kDa) in EndoC-βH1 cells transfected with a synthetic inhibitor of hsa-miR-184-3p. Data are shown as mean±SD of fold change values of normalized CRTC1/β-Act ratio of miR-184-p inhibitor vs miR-Ctr inhibitor transfected samples; statistics using paired Student's t-test (n=6).

Overall, these data suggest that miR-184-3p directly regulates the expression of CRTC1 in human β-cell line EndoC-βH1 as well as in other cellular contexts.

### CRTC1 upregulation protects β-cells from lipotoxic- or pro-inflammatory-mediated apoptosis

In order to verify the contribution and function of CRTC1 to miR-184-3p mediated protection effects from apoptosis, we directly overexpressed CRTC1 in MIN6 cells and then exposed them to palmitate or to a mix of pro-inflammatory cytokines. The results showed that CRTC1 overexpression (**Fig. 4a**) induced β-cell protection from palmitate exposure as shown by active Caspase-3 staining (**Fig. 4b and Suppl. Fig. 3a)**and by pyknotic nuclei count (**Suppl. Fig.3b**). Furthermore, CRTC1 overexpression (**Fig. 4c**) was able to protect MIN6 from cytokines-induced apoptosis as well (**Fig. 4d and Suppl. Fig. 3c, Suppl. Fig. 3d)**.

**Figure 4.**
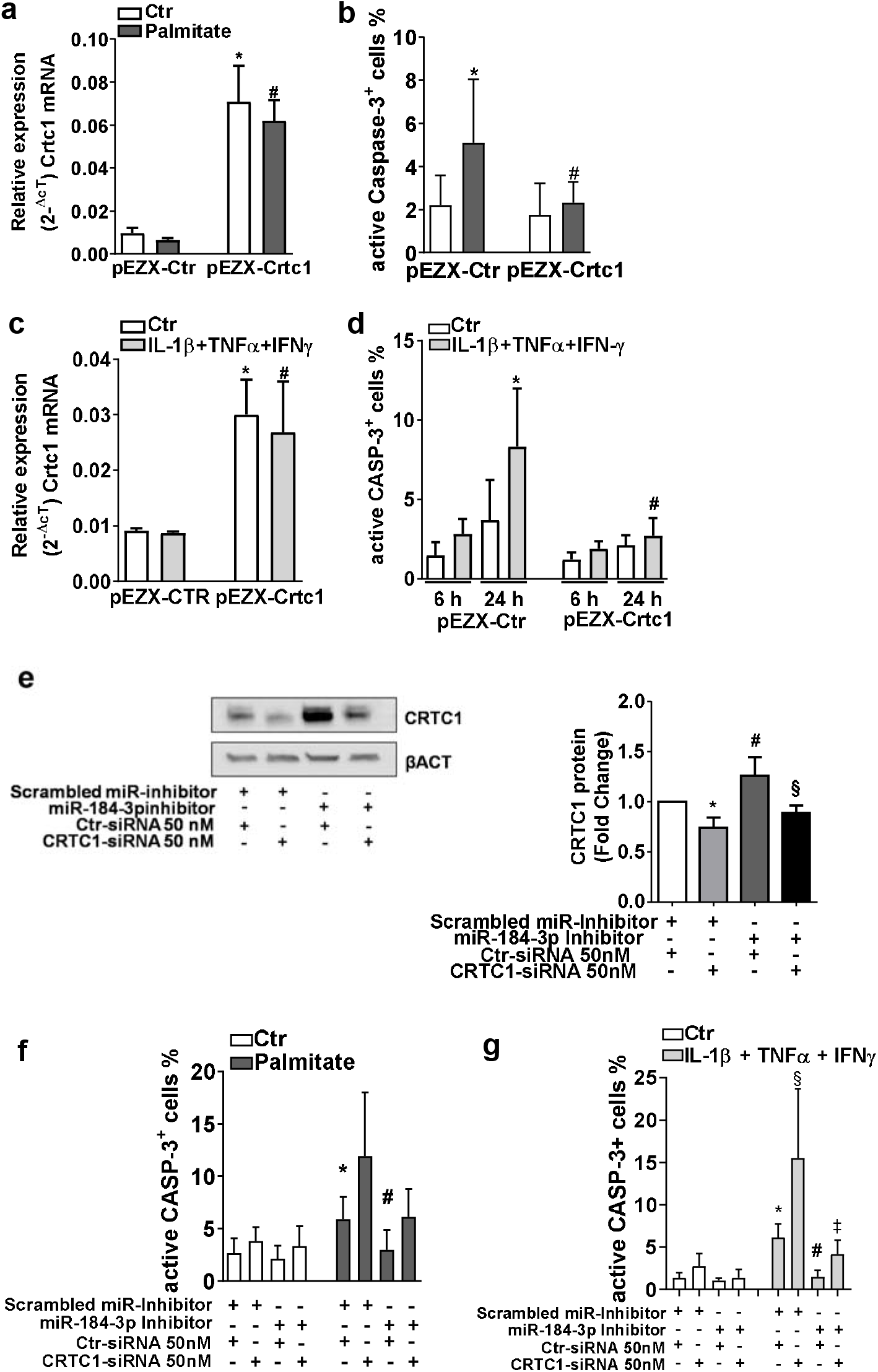
miR1-184-3p-mediated protection from palmitate and cytokines-induced apoptosis is dependent on CRTC1 upregulation. **(a)(c)** RT-Real Time PCR of CRTC1 mRNA to validate its overexpression upon specific plasmid vector transfection in MIN6 subjected to lipotoxic (a) or inflammatory stress (c); data are shown as mean ± SD of 2^-ΔCt^ relative expression normalized to β-act and Gapdh.* p≤0.05 vs pEZX-Ctr transfected using Wilcoxon test (n=4 independent experiments). **(b)(d)** Cytofluorimetric assessment of active Caspase-3+ (active-CASP-3+) MIN6 cells overexpressing CRTC1 after 24h of palmitate treatment (b) (*p<0.0001 vs not treated pEZX-Ctr transfected; #p<0.0001 vs palmitate treated pEZX-Ctr transfected) or cytokines mix (IL-1β+TNF-α+IFN-γ) treatment for 6h and 24h (d) (*p=0.0266 vs 24h not treated pEZX-Ctr transfected; #p=0.0359 vs 24h cytokines treated pEZX-Ctr transfected; values are reported as mean percentage ± SD of Caspase-3+ cells on total live cells; statistics performed using one-way ANOVA with Bonferroni’s multiple comparison test (n=6 independent experiments). **(e-*left panel***) Western blot analysis of CRTC1 (78 kDa) and housekeeping control β-ACT (43 kDa) in EndoC-βH1 cells transfected with CRTC1 siRNA (50 nM) in combination or not with miR-184-3p inhibitor; (**e-*right panel***) densitometric analysis of CRTC1 protein expression upon β-ACT normalization (*p=0.044 vs Scrambled miR-inhibitor + Ctr siRNA transfected; #p=0.0409 vs Scrambled miR-inhibitor + Ctr siRNA transfected; § p=0.0002 vs scrambled miR-inhibitor + CRTC1 siRNA transfected). Data are shown as mean ± SD of fold change values of normalized CRTC1/β-ACT ratio; statistics performed using one-way ANOVA with Bonferroni’s multiple comparison test (n=6 independent experiments). **(f),(g)** Cytofluorimetric assessment of Caspase-3^+^ EndoC-βH1 cells transfected with CRTC1 siRNA (50nM) in combination or not with the synthetic inhibitor of hsa-miR-184-3p and respectively treated with palmitate for 24h (f) (*p=0.02 vs not treated and Scrambled miR-inhibitor + Ctr siRNA transfected; #p<0.0001 vs not treated miR-184 inhibitor + Ctr siRNA transfected) or treated with cytokines for 24h (g) (*p<0.0001 vs not treated Scrambled miR inhibitor + Ctr siRNA transfected; § vs not treated Scrambled miR-inhibitor + CRTC1 siRNA; #p=0.0005 vs not treated and miR-184 inhibitor + Ctr siRNA transfected; ‡ p<0.0001 vs not treated miR-184-3p inhibitor + CRTC1 siRNA. Statistics performed using one-way ANOVA with Bonferroni’s multiple comparison test (n=6 independent experiments).

Importantly, the direct contribution of CRTC1 to miR-184-3p mediated protection was confirmed in the human context using EndoC-βH1 cells. Indeed, in order to verify whether CRTC1 had a role in β-cell protection mediated by miR-184-3p expression levels reduction, CRTC1 was knocked down, alone or in combination with miR-184-3p inhibition, during palmitate or cytokines treatment. The expression of CRTC1 was verified by western blot analysis (**Fig. 4e**). In addition, in order to further confirm CRTC1 protein expression modulation upon siRNA or miR-184-3p inhibition, we adopted a targeted mass spectrometry (MS) quantitation approach which confirmed western blot data (**Suppl. Fig. 4a and Suppl. Fig. 4b**); the results showed the expected expression pattern which was in line with CRTC1 silencing using specific siRNA, its upregulation upon miR-184-3p inhibition and no significant differential expression vs control by transfecting CRTC1 siRNA alongside with miR-184-3p inhibitor (**Fig. 4e- left and right panels**). As expected, the apoptosis rate, measured by active Caspase-3 staining, was reduced upon miR-184-3p inhibition both in palmitate and in cytokines-exposed EndoC-βH1 cells. Interestingly, CRTC1 silencing significantly increased apoptosis rate upon palmitate stress-induction and, importantly, completely abrogated the protection effects exerted by miR-184-3p inhibition (**Fig. 4f**) (**Supplementary Figure 5a**/**b**). A similar pattern was observed also in cytokines-treated EndoC-βH1 cells (**Fig. 4g**) (**Supplementary Figure 6a**-**6b**). Collectively, these data demonstrate the central role of CRTC1 in apoptosis protection mechanism exerted by miR-184-3p inhibition in β-cells, both during lipotoxic stress and pro-inflammatory conditions.

### β-cell specific transcription factor NKX6.1 regulates miR-184-3p expression

Although miR-184-3p downstream mechanisms have been previously partially addressed and further elucidated in the present work, upstream factors determining miRNA downregulation need to be further explored. Therefore, in order to shed light onto the regulatory cues modulating miR-184 gene transcriptional activity, we analyzed miR-184 promoter sequence, taking into account the 0,5 kb upstream sequence (**Suppl. Figure 7**) previously indicated as miR-184 gene proximal promoter regulatory region ^38^. *In-silico* analysis of human miR-184 promoter showed 234 predicted TFBS. Among them, we identified 3 binding sites for the β-cell specific transcription factor NKX6.1, while only a single binding site was detected in murine miR-184-3p promoter sequence (**Fig. 5a-left and right panel and Suppl. File 1**). NKX6.1 DNA Responsive Elements (DRE) were characterized by the typical homeodomain core sequence binding motif belonging to NKX6.1 (NNTTAANN)^41^ (**Fig 5a- left panel**). Given the fundamental role of NKX6.1 in endocrine pancreatic cell differentiation and in the maintenance of β-cell functional identity^42^, we focused on its putative regulation of miR-184 gene transcription. In order to investigate whether NKX6.1 actually regulates the expression of miR-184, we measured its expression in pancreatic islets isolated from n=10 non-diabetic and n=6 T2D donors (**Supplementary Table 1**). The results showed a significant reduction of NKX6.1 in T2D vs. non-diabetic pancreatic islets (**Fig. 5b**), in line with the downregulation of miR-184-3p. Furthermore, the expression levels of NKX6.1 resulted positively correlated to miR-184-3p (**Fig. 5c**). Additionally, in order to evaluate the role of NKX6.1 in miR-184 transcription, we silenced it in EndoC-βH1 cells (**Fig. 5d and Fig. 5e)** and then measured miR-184-3p expression; the results showed a significant reduction of miR-184-3p upon NKX6.1 siRNA transfection (**Fig. 5f**), thus demonstrating a direct or indirect link between them.

**Figure 5.**
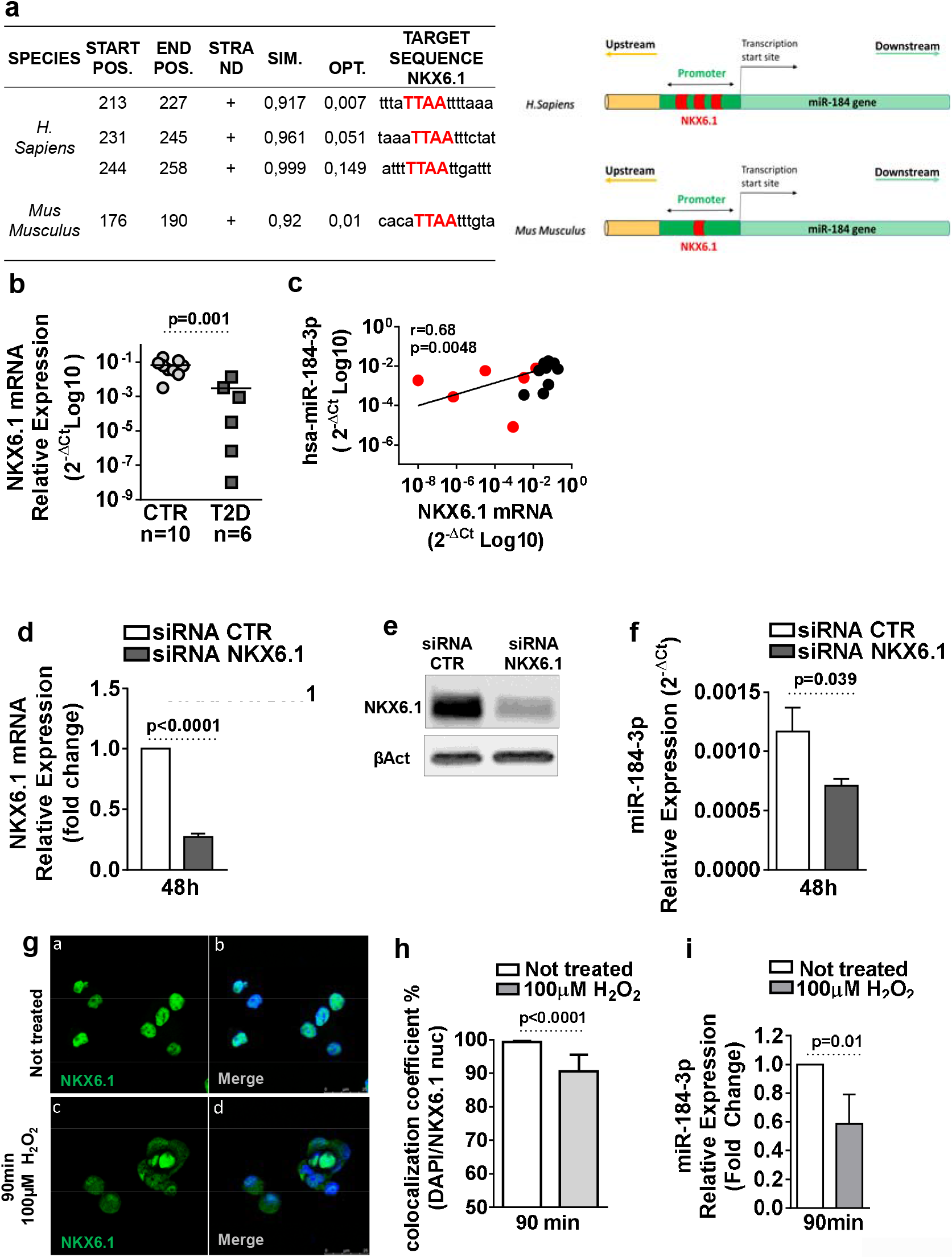
NKX6.1 regulates human and murine miR-184-3p expression. **(a-*left panel*)** Table reporting predicted binding sites of NKX6.1 in human and mouse miR-184 gene promoter using MatInspector. For each predicted binding site the following parameters are reported: species, start/end nucleotide (Start Pos, End Pos) of TF matrix (respect to −500bp from miR-184 TSS, see sequence in Supplementary Material), sequence strand, Matrix similarity (SIM.), optimized matrix similarity threshold (OPT.) and target sequence (see Supplementary file 1). (**a-*right panel***) representative scheme reporting human and mouse MIR184 gene promoter and NKX6.1 binding sites (in red). **(b)** RT-Real-Time PCR expression analysis of NKX6.1 in T2D human pancreatic islets (n=6; dark grey squares) vs non-diabetic donors (n=10; light grey circles); data are shown as 2^-ΔCt^ relative expression; statistics using Mann Whitney U test. **(c)** Correlation analysis between NKX6.1 and miR-184-3p expression in human pancreatic islets from T2D and non-diabetic donors (n=16). Values are reported as 2^-ΔCt (Log10)^ relative expression; p-value and r-value by Spearman R correlation test. **(d, e)** 48h silencing of NKX6.1 using siRNA (25 nM) in EndoC-βH1 cell line showed mRNA (d) and protein (e) expression reduction; statistics using Mann Whitney U test (n=4) **(f)** RT-Real Time PCR of miR-184-3p expression in EndoC-βH1 cells upon 48h NKX6.1 siRNA transfection; data are reported as mean±SD of 2^-ΔCt^ normalized values. Statistics performed using Mann Whitney U test (n=4 independent experiments). **(g)** Immunofluorescence analysis of Nkx6.1 (green) and nuclei (blue) in MIN6 cells treated or not with 100μM H_2_O_2_; panel a (Nkx6.1) and panel b (merged signal: Nkx6.1/nuclei) showing not treated MIN6 cells; panel c (Nkx6.1) and panel d (merged signal: Nkx6.1/nuclei) showing H_2_O_2_ treated MIN6 cells; scale bar 25um. **(h)** Colocalization coefficient quantification analysis between Nkx6.1 and nuclei staining with or without treatment with 100μM H_2_O_2_. Data are reported as mean ± SD of the percentage of Nkx6.1 signal overlapping nuclei staining. statistics using Mann Whitney U test (n=5). **(i)** RT-Real Time PCR analysis of miR-184-3p expression in MIN6 cells upon 100μM H_2_O_2_ treatment. Data are shown as mean ± SD of fold change of H_2_O_2_ treated MIN6 cells vs not treated samples; statistics using Mann Whitney U test (n=5).

Next, we sought to determine whether NKX6.1 has a role in miR-184-3p expression in murine β-cells as well. Therefore, we evaluated whether the translocation of NKX6.1 from nucleus to cytoplasm could be sufficient to induce a reduction of miR-184-3p transcription. As a matter of fact, it was previously demonstrated that murine β-cell lines exposed to oxidative stress showed the translocation of NKX6.1 from nucleus to cytoplasm as a mechanism of β-cell dysfunction^43^. Therefore, we exposed MIN6 cells to H_2_O_2_ stimuli for 90 min and then evaluated the translocation of NKX6.1 and the expression of miR-184-3p. In MIN6 cultured in basal condition, NKX6.1 was mainly present in the nucleus as a sign of its functional and transcriptional activity (**Fig. 5g, panel a-b**). Upon H_2_O_2_ treatment, NKX6.1 was partially translocated to cytoplasm, as shown by immunofluorescence staining (**Fig. 5g, panel c-d)** and by NKX6.1/nuclei colocalization analysis **(Fig. 5g**). Interestingly, NKX6.1 nucleus-cytoplasm shuttling was accompanied by a significant reduction of miR-184-3p expression (**Fig. 5h**) as well as of MafA (**Supplementary Fig. 8)**, a previously demonstrated and well known target gene of NKX6.1^42,44^.

### NKX6.1 directly binds miR-184 promoter sequence in human and murine β-cells

With the purpose to distinguish between a direct regulation of NKX6.1 on miR-184 promoter or an indirect role through other putative targeting factors, we performed a ChIP analysis of human miR-184 proximal promoter regulatory region, containing NKX6.1 core binding sites. NKX6.1 efficiently binds to the putative selected sequence region of miR-184 promoter in EndoC-βH1 cells (**Fig. 6a**) and, furthermore, its binding was significantly reduced upon NKX6.1 siRNA transfection vs scrambled siRNA control (**Fig. 6a**). Additionally, immunoprecipitated chromatin related to miR-184-3p promoter sequence also showed a reduced acetylation on H3K9 and H3K14, thus suggesting a lower site specific chromatin activation state in the absence of NKX6.1 (**Fig.6b**).

**Figure 6.**
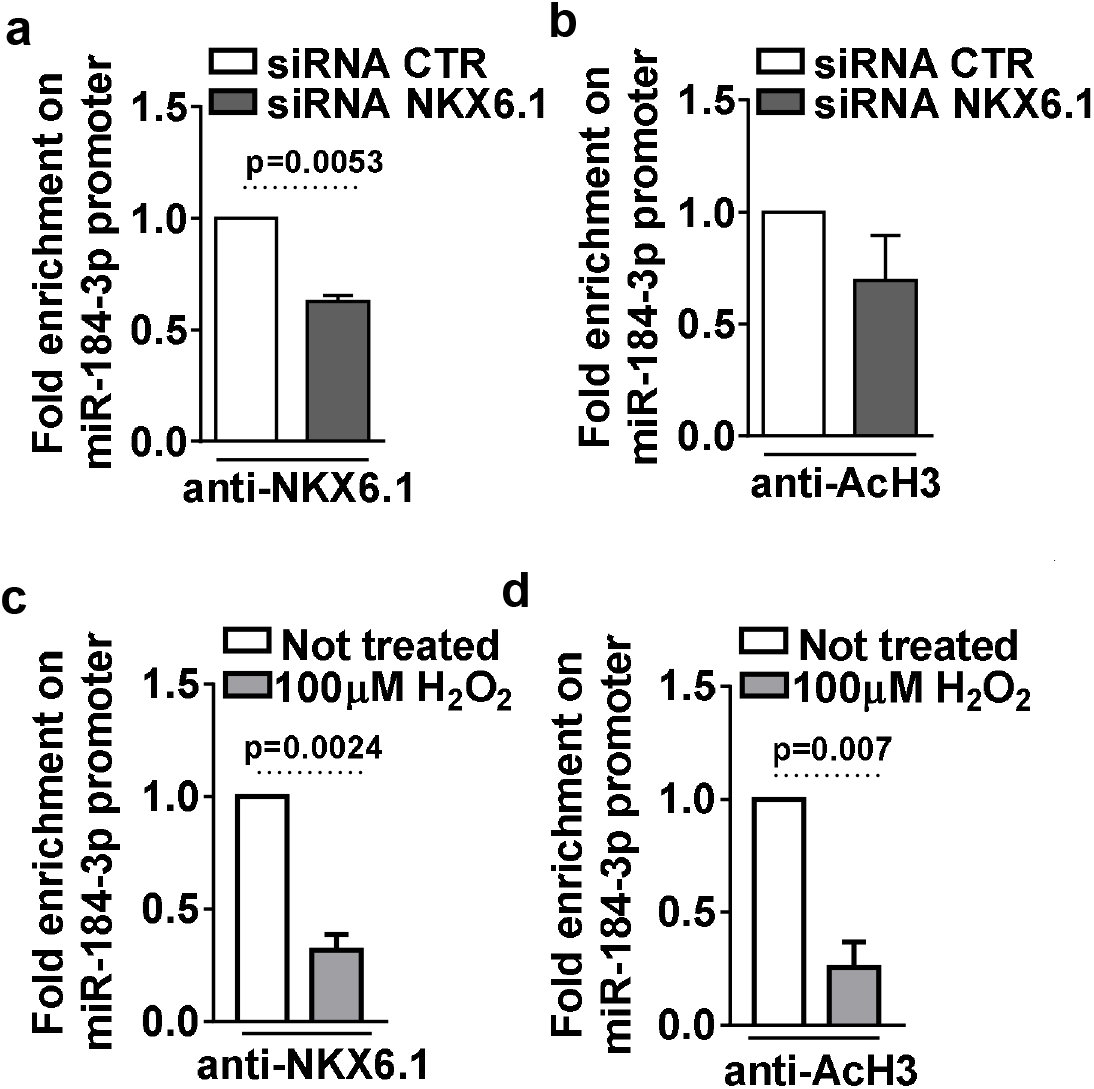
NKX6-1 directly binds to miR-184 promoter. (**a,b**) ChIP-qPCR of NKX6.1 binding on human miR-184 promoter selected sequence; data are shown as fold change of miR-184 promoter sequence bound by anti-NKX6.1 antibody vs negative control isotypic IgG. ChIP-qPCR of NKX6.1 binding (a) or Histone H3 acetylation (AcH3) (b) on miR-184 promoter, with or without NKX6.1 siRNA and normalized to IgG control; values are reported as fold change of NKX6.1 siRNA transfected EndoC-βH1 cells vs scrambled siRNA; statistics using Mann Whitney U test (n=4-5). **(c,d)** ChIP-Real Time PCR of NKX6.1 binding on murine miR-184 promoter selected sequence; data are shown as fold change of anti-NKX6.1 antibody vs negative control isotypic IgG; statistics using Mann Whitney U test, n=4. ChIP-qPCR of NKX6.1 binding (c) or Histone H3 acetylation (AcH3) (d) on murine miR-184 promoter, with or without 100μM H_2_O_2_ treatment in MIN6 cells; values are reported as fold change of 100μM H_2_O_2_ treated samples vs not-treated; statistics using Mann Whitney U test (n=4).

ChIP-qPCR analysis revealed that NKX6.1 actually binds to the promoter regulatory region upstream miR-184 TSS also in MIN6 cells (**Fig. 6c-6d**). Of note, oxidative stress induced NKX6.1 translocation (**Fig.5 g-h**), reduced the binding to miR-184 promoter region (**Fig. 6c**) and lowered the acetylation of H3K9 and H3K14 (**Fig. 6d**) as a result of the decrease of the chromatin activation state.

Overall, these data demonstrate that NKX6.1 is part of the transcriptional machinery leading to the regulation of miR-184-3p expression both in human and in murine β-cells, therefore linking miR-184-3p reduction to the potential alteration of NKX6.1 expression and subcellular localization in type 2 diabetes.

## Discussion

The present study provides evidence of a novel mechanism driven by miR-184-3p in β-cells, partially regulated by NKX6.1 and which supports a role for this miRNA in T2D pathophysiology. Importantly, our results are corroborated by previously reported findings, including miR-184-3p expression reduction in T2D human pancreatic islets^20^ and in those derived from db/db and ob/ob murine models^20,21^. In addition, we further demonstrated the role of miR-184-3p in β-cell protection from metabolic or inflammatory-mediated apoptosis ^21^.

MiR-184-3p functions have been already partially addressed in previous studies which shed light onto the importance of this miRNA in β-cells physiology and in the pathogenesis of T2D; such studies revealed that: (*i*) miR-184-3p has been shown to be one of the β-cell/pancreatic islet enriched miRNAs^20^; (*ii*) it regulates insulin secretion by repressing the expression of SLC25A22 gene^23^ and regulates compensatory β-cell expansion by targeting AGO2^20^; (*iv*) it is partially regulated by the energy sensor AMPK^24^.

Despite this bulk of information, none of these studies deeply characterized the molecular factors mediating the previously observed anti-apoptotic mechanism residing downstream its reduction; nonetheless, the evidence of upstream regulators of miR-184-3p expression are scarce. Here, by using downstream and upstream bioinformatic approaches followed by the analysis of T2D and non-diabetic human pancreatic islets, the human β-cell lines EndoC-βH1 and 1.1B4 as well as of MIN6 murine β-cells, we identified both targets of miR-184-3p and putative transcription factors which may mediate its transcriptional regulation. Indeed, we identified and validated CRTC1 as a downstream target of miR-184-3p. CRTC1 is a member of the cAMP-regulated transcriptional co-activators (CRTCs) family which binds to the bZIP domain of CREB transcription factor, leading to an increased occupancy over the DNA CRE-responsive elements, thus activating the expression of a specific pattern of genes^45,46,47^. As a matter of fact, CRTC1 is required for the efficient induction of CREB target genes. Additionally, CRTC1 activates CRE-responsive elements even in the absence of extracellular signals, thus making CRTC1 a master modulator of CREB activity^48^. Here, we showed that CRTC1 overexpression in β-cells, directly protects them from lipotoxic-mediated apoptosis. The protection effect exerted by CRTC1 axis covers also inflammatory stress (IL-1β+TNF-α+IFN-γ), indicating that such protection process represents a basic β-cell survival mechanism. Interestingly, in human β-cells, miR-184-3p mediated-protection is totally dependent on CRTC1 overexpression, being completely abrogated when CRTC1 is silenced. These results are in line with the previously observed role of CREB-CRTC; indeed, transgenic mice expressing a defective form of CREB (A-CREB) developed hyperglycaemia, due to an increased β-cell apoptosis rate^49–51^. Therefore, it is conceivable that the reduction or the overexpression of its transcriptional coactivator CRTC1, which modulates CREB activity and, in turn, the expression of its target genes, may contribute to the exacerbation or protection from apoptosis, respectively. Moreover, this model fits with the mechanism described by Martinez-Sanchez et al. who observed a circuitry between AMPK-miR-184-3p and β-cell survival^24^. Indeed, they reported that the reduction of AMPK activity due to high glucose stimuli^52^ induces a reduction of miR-184-3p expression; we can hypothesize that this phenomena is paralleled by high glucose-dependent increase of cAMP which, in turn, elicits further activation of CRTC1^53^ whose expression is enhanced by miR-184-3p inhibition.

While the protection effect exerted by CRTC1 was conserved between human and mouse β-cells, the mechanism leading to its overexpression and involving miR-184-3p seems not to be conserved as well. Indeed, murine Crtc1 is not a predicted target of miR-184-3p neither its expression was modulated upon miR-184-3p inhibition (data not shown). However, the protection function exerted by miR-184-3p downregulation was observed both in human and mouse; we speculate that additional mechanisms not involving CRTC1 in mouse β-cells may mediate the protection effect upon miR-184-3p inhibition.

An additional unanswered point resides in the mechanism through which CRTC1 overexpression protects β-cells from apoptosis; indeed, the pattern of downstream activated target genes involved in such process remains to be further investigated.

A potential role in the protection from apoptosis may be played by miR-212/miR-132 axis, whose expression in β-cells was already demonstrated to be controlled by CRTC1-CREB^54^ complex and whose target genes are involved in the protection from apoptosis in multiple cell types including β-cells^55,56^. Collectively, these results open to a potential β-cell protective/survival cue during T2D, exerted by miR-184-3p-CRTC1 axis and leading to protection from apoptosis. This view is in line with recent findings indicating that β-cell death may contribute to a lesser extent to the reduction of functional β-cell mass than previously noted and that other mechanisms are involved in β-cell dysfunction^4,5,57^. Our study provides evidence that NKX6.1 is downregulated in pancreatic islets of T2D donors and, more importantly, that it regulates the expression of miR-184-3p both in human and mouse β-cells by specifically binding to DNA responsive elements present within its proximal promoter. NKX6.1 has been demonstrated to be a β-cell enriched transcription factor which controls a defined genes network necessary for specific β-cell function^43^; additionally, NKX6.1 maintains phenotype identity by controlling some β-cell specific traits^44,58^. Of importance, previous studies also attributed a critical role to NKX6.1 in β-cell dedifferentiation. Indeed, several evidences indicated that: *i*) NKX6.1 translocates from nucleus to cytoplasm in β-cells of T2D donors and db/db mice, and was accompanied by the activation of aldehyde dehydrogenase-1 a3 (Aldh1a3) and concomitant activation of typical undifferentiated endocrine cells marker genes^59,60^; *ii*) loss of function of NKX6.1 has been linked to the loss of β-cell specific traits causing the disruption of some specific gene-regulatory networks, thus leading to the loss of β-cell identity^22^. Our data confirmed the reduced expression of NKX6.1 in pancreatic islets of T2D donors vs non-diabetic controls; moreover, NKX6.1 islets expression was positively correlated with the expression of miR-184-3p in consequence of a direct transcriptional regulation. Although additional studies are needed to address the exact contribution of NKX6.1 reduced expression and its nucleus to cytoplasm translocation to miR-184-3p expression, we can speculate that both these processes can occur in stress conditions during T2D ^59^ thus leading to miR-184-3p downregulation. Indeed, the direct link between NKX6.1 loss of function and β-cell dedifferentiation has been reported in several previous studies^5,59^; such evidence may suggest that in consequence of NKX6.1 functional reduction due to a dedifferentiation process, miR-184-3p expression can be decreased, thus leading to CRTC1 upregulation and to β-cell protection. Therefore, although dysfunctional in consequence of a dedifferentiation process, β-cells can be partially protected from apoptosis, thus indicating a potential association between dedifferentiation and protection. Such model is corroborated by several recent findings correlating β-cell dedifferentiated phenotype to a protection from both metabolic and inflammatory insults^6,61,62^.

In conclusion, we propose a model (**Fig. 7**) where the observed reduction of miR-184-3p and the consequent upregulation of its target gene CRTC1 in pancreatic islets of T2D donors follow the reduction/translocation of NKX6.1 observed in de-differentiating and/or dysfunctional β-cells in type 2 diabetes, thus opening to novel potential possibilities of therapeutic interventions.

**Figure 7.**
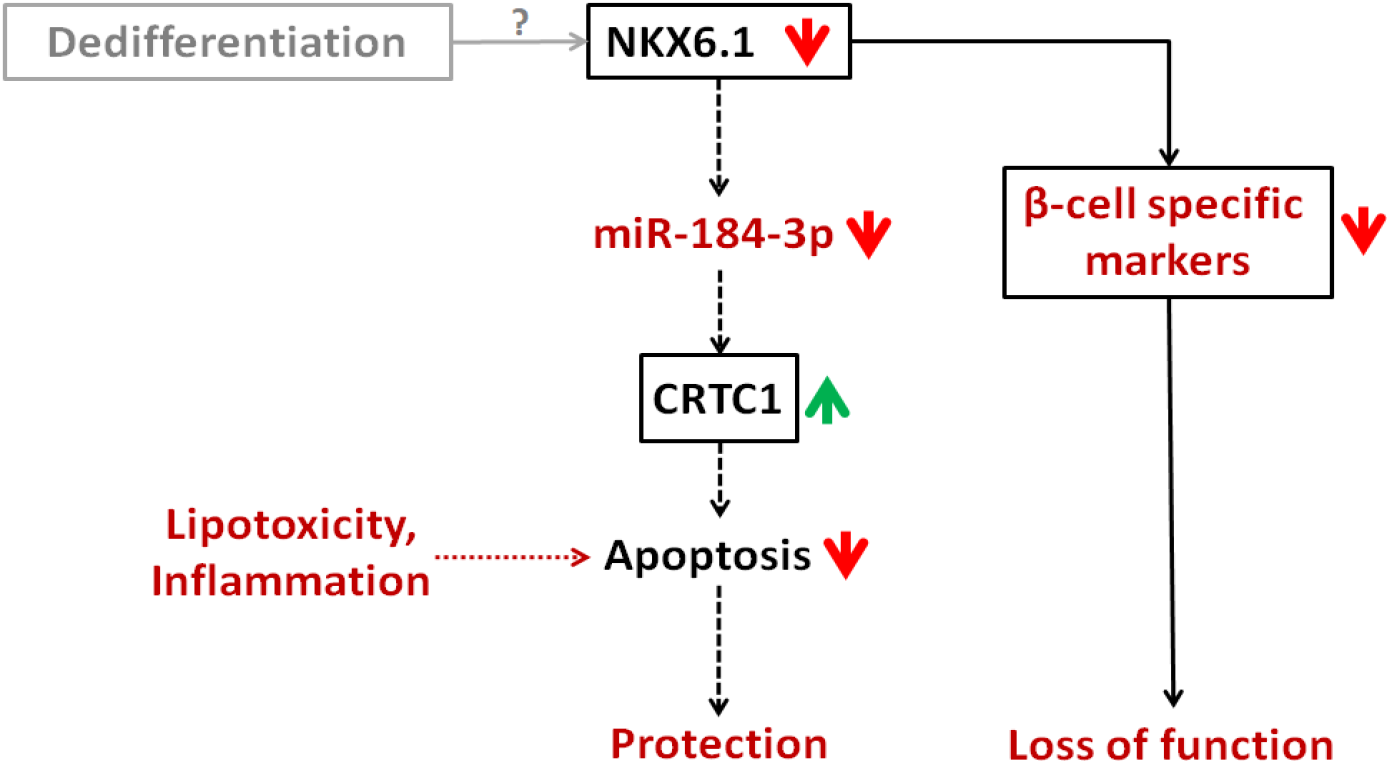
Working model of upstream and downstream miR-184-3p molecular mechanisms in β-cells protection.

## Acknowledgements

The Secretarial help of Maddalena Prencipe and Alessandra Mechini was greatly appreciated.

