## Supplementary Material for "Reduced miR-184-3p expression occurring in Type 2 diabetic pancreatic islets protects β-cells from lipotoxic and proinflammatory apoptosis via a CRTC1-dependent mechanism"

**Supplementary Table 1**. **Main characteristics of T2D and non diabetic multiorgan donors**. Sex, Age (years), BMI (Kg/m^2^), disease duration (years), ICU (Intensive Care Unit) stay (days) and cause of death are reported. The analyses performed on each donor samples are reported as well (miR-184-3p, CRTC1 and/or NKX6.1 expression).

|  | **Samples** | **Sex (M/F)** | **Age (years)** | **(BMI) [Kg/m2]** | **Disease**  **Duration (years)** | **ICU stay (days)** | **Cause of Death** | **miR-184-3p analysis** | **CRTC1 analysis** | **NKX6.1 expression** |
| --- | --- | --- | --- | --- | --- | --- | --- | --- | --- | --- |
| **No diabetes** | Hi1 CTR | F | 78 | -- | n/a | 2 | Cardiovasc | yes | no | no |
|  | Hi2 CTR | M | 54 | 23,1 | n/a | 6 | Cardiovasc | yes | no | no |
|  | Hi3 CTR | F | 59 | 27,7 | n/a | 12 | Cardiovasc | yes | no | no |
|  | Hi4 CTR | M | 53 | 34,6 | n/a | -- | -- | yes | yes | yes |
|  | Hi5 CTR | M | 52 | 34,2 | n/a | -- | -- | yes | yes | yes |
|  | Hi6 CTR | M | 50 | 27,4 | n/a | -- | -- | yes | yes | yes |
|  | Hi7 CTR | M | 55 | 28 | n/a | -- | -- | yes | yes | yes |
|  | Hi8 CTR | F | 79 | 23,9 | n/a | 1 | Cardiovasc | yes | yes | yes |
|  | Hi9 CTR | M | 59 | 26,7 | n/a | -- | Cardiovasc | yes | yes | yes |
|  | Hi10 CTR | F | 74 | 23,4 | n/a | 9 | Cardiovasc | yes | yes | yes |
|  | Hi11 CTR | F | 53 | 25,7 | n/a | 1 | -- | yes | yes | yes |
|  | Hi12 CTR | F | 81 | 25,9 | n/a | -- | Cardiovasc | yes | yes | yes |
|  | Hi13 CTR | M | 74 | 24,8 | n/a | 5 | Cardiovasc | yes | yes | yes |
|  | Hi1 T2D | M | 76 | 26 | -- | 2 | Cardiovasc | yes | yes | yes |
| **Type 2 diabetes** | Hi2 T2D | F | 54 | 23,9 | 9 | 8 | Cardiovasc | yes | no | no |
|  | Hi3 T2D | F | 73 | 29,3 | 8 | 4 | Cardiovasc | yes | no | no |
|  | Hi4 T2D | M | 78 | 25,9 | 25 | 2 | Trauma | yes | yes | yes |
|  | Hi5 T2D | M | 69 | 27,7 | 5 | 5 | Trauma | yes | yes | yes |
|  | Hi6 T2D | M | 72 | -- | -- | -- | Cardiovasc | yes | no | no |
|  | Hi7 T2D | M | 78 | 26,1 | -- | -- | Cardiovasc | yes | yes | yes |
|  | Hi8 T2D | M | 69 | 25,8 | 4 | 1 | Cardiovasc | yes | yes | yes |
|  | Hi9 T2D | F | 81 | 23,4 | 18 | 1 | Cardiovasc | yes | yes | yes |

| **Target gene** | | **Representative transcript** | | **Gene name** | | **Total Sites** | | **Cumulative Weighted Score++** |
| --- | --- | --- | --- | --- | --- | --- | --- | --- |
| SF1 | | ENST00000377390.3 | | splicing factor 1 | | 3 | | -0.79 |
| ***CRTC1*** | | ***ENST00000338797.6*** | | ***CREB regulated transcription coactivator 1*** | | ***3*** | | ***-0.73*** |
| CRISPLD2 | | ENST00000262424.5 | | cysteine-rich secretory protein LCCL domain containing 2 | | 3 | | -0.29 |
| EPB41L5 | | ENST00000443902.2 | | erythrocyte membrane protein band 4.1 like 5 | | 2 | | -0.93 |
| ALDH4A1 | | ENST00000375341.3 | | aldehyde dehydrogenase 4 family, member A1 | | 2 | | -0.72 |
| SLC25A22 | | | ENST00000320230.5 | solute carrier family 25 (mitochondrial carrier: glutamate), member 22 | 2 | | -0.60 | |
| MFRP | ENST00000555262.1 | | | membrane frizzled-related protein | | 2 | | -0.47 |
| NFIC | ENST00000589123.1 | | | nuclear factor I/C (CCAAT-binding transcription factor) | | 2 | | -0.30 |
| SIDT2 | ENST00000324225.4 | | | SID1 transmembrane family, member 2 | | 1 | | -0.68 |
| NUS1 | ENST00000368494.3 | | | nuclear undecaprenyl pyrophosphate synthase 1 homolog (S. cerevisiae) | | 1 | | -0.67 |
| FBXO28 | ENST00000424254.2 | | | F-box protein 28 | | 1 | | -0.60 |
| ZNF865 | ENST00000568956.1 | | | zinc finger protein 865 | | 1 | | -0.60 |
| HAND2 | ENST00000359562.4 | | | heart and neural crest derivatives expressed 2 | | 1 | | -0.60 |
| ZNF740 | ENST00000416904.3 | | | zinc finger protein 740 | | 1 | | -0.59 |
| PPAP2B | ENST00000371250.3 | | | phosphatidic acid phosphatase type 2B | | 1 | | -0.48 |
| REEP5 | ENST00000545426.1 | | | receptor accessory protein 5 | | 1 | | -0.47 |
| STC2 | ENST00000265087.4 | | | stanniocalcin 2 | | 1 | | -0.46 |
| NCOR2 | ENST00000405201.1 | | | nuclear receptor corepressor 2 | | 1 | | -0.43 |
| C15orf37 | ENST00000560255.1 | | | chromosome 15 open reading frame 37 | | 1 | | -0.33 |
| FCRLB | ENST00000336830.5 | | | Fc receptor-like B | | 1 | | -0.31 |
| CARM1 | ENST00000327064.4 | | | coactivator-associated arginine methyltransferase 1 | | 1 | | -0.30 |
| MLEC | ENST00000228506.3 | | | malectin | | 1 | | -0.29 |
| HIF1AN | ENST00000299163.6 | | | hypoxia inducible factor 1, alpha subunit inhibitor | | 1 | | -0.27 |
| TNRC6B | ENST00000335727.9 | | | trinucleotide repeat containing 6B | | 1 | | -0.24 |
| CBX6 | ENST00000407418.3 | | | chromobox homolog 6 | | 1 | | -0.21 |
| C20orf112 | ENST00000359676.5 | | | chromosome 20 open reading frame 112 | | 1 | | -0.21 |
| ZIC4 | ENST00000383075.3 | | | Zic family member 4 | | 1 | | -0.17 |
| AGO2 | ENST00000220592.5 | | | argonaute RISC catalytic component 2 | | 1 | | -0.12 |
| DBNL | ENST00000494774.1 | | | drebrin-like | | 1 | | 0 |

**Supplementary Table 2. List of miR-184-3p predicted target genes**. Conserved and high ranked miR-184-3p predicted target genes by Targetscan7.1 algorithm. Target gene name, Ensembl transcript ID, extended gene name, total number of sites in 3’UTR and Cumulative Weighted Score are reported. Target genes are listed according to the number of miRNA sites in the 3’UTR.

**Supplementary Table 3. List of primers used for RT Real-Time PCR.** List of Taqman primers used to evaluate genes and microRNAs expression.

| **Gene name** | **ID code** | **Species** |
| --- | --- | --- |
| ACTIN β | 4333762F | Human |
| GAPDH | 4333764F | Human |
| CRTC1 | Hs00993064_m1 | Human |
| NKX6.1 | Hs00232355_m1 | Human |
| ACTIN β | Mm02619580_g1 | Mouse |
| GAPDH | Mm99999915_g1 | Mouse |
| CRTC1 | Mm01349190_m1 | Mouse |
| NKX6.1 | Mm00454961_m1 | Mouse |
| MAFA | Mm00845206_s1 | Mouse |

| **microRNA name** | **ID code** | **Species** |
| --- | --- | --- |
| U6 snRNA | 001973 | Human & Mouse |
| dme-miR-7 | 000268 | Human & Mouse |
| miR-184 | 000485 | Human & Mouse |

**Supplementary Figure legends**

**Supplementary Figure 1. Representative cytofluorimetric plots and pyknotic nuclei count upon palmitate or cytokines treatment and miR-184-3p inhibition in EndoC-βH1 cells. (a)(c)** Representative cytofluorimetric plot of EndoC-βH1 cells transfected with a synthetic inhibitor of miR-184-3p or with a CTR inhibitor and treated with palmitate or cytokines**.** (**b**) (**d**) Pycnotic nuclei count of EndoC-βH1 cells transfected with a synthetic inhibitor of miR-184-3p and treated with palmitate or cytokines mix. Data are reported as mean percentage ±SD of pyknotic nuclei cells on total nuclei. * p≤0.05 vs not-treated miR-CTR inhibitor transfected; # p≤0.05 (vs cytokines treated + miR-CTR inhibitor transfected). Statistics performed using ANOVA analysis with Bonferroni’s multiple comparison test (n=4 independent experiments).

**Supplementary Figure 2. CRTC1 mRNA and protein expression analysis upon miR-184 inhibition in 1.1B4 human pancreatic cell line and in HeLa cells. (a,c)** RT Real-Time PCR of CRTC1 expression upon miR-184-3p inhibition respectively in 1.1B4 (a) (48h) and in HeLa cell (c) (24h). Data are reported as mean±SD of fold change expression relative to control. (**b,d**) CRTC1 Western Blot analysis upon miR-184-3p inhibition respectively in 1.1B4 (b) (48h) and in HeLa cells (d) (24h). Values are reported as mean±SD of fold change relative to control of CRTC1/β-ACT ratio. Statistics performed using paired Student's t-test (*p≤0.05, n=4 independent experiments).

**Supplementary Fig. 3. Representative cytofluorimetric plots and pyknotic nuclei count after palmitate or cytokines mix treatment and CRTC1 overexpression in MIN6 cells. (a,c)** Representative cytofluorimetric plots of MIN6 cells treated with palmitate (a) or cytokines mix (c) (24h treatment cytofluorimetric plots showed) and transfected or not with CRTC1 overexpressing vector**.** (**b**) Pyknotic nuclei count of MIN6 cells treated with palmitate (48h) or cytokines mix (6h or 24h) and transfected or not with CRTC1 overexpressing vector. Data are reported as mean±SD of the percentage of positive pyknotic nuclei cells on total nuclei.*p≤0.05 (vs not treated+ pEZX-Ctr tranfected), #p≤0.05(vs palm. treated + pEZX-Ctr transfected), statistics using ANOVA analysis with Bonferroni’s multiple comparison test (n=4). (**d**) Pyknotic nuclei count of MIN6 cells treated with cytokines mix (6h or 24h) and transfected or not with CRTC1 overexpressing vector. Data are reported as mean±SD of the percentage of positive pyknotic nuclei cells on total nuclei.*p≤0.05 (vs not treated + pEZX-Ctr tranfected, 24h), #p≤0.05 (vs cytokines treated + pEZX-Ctr transfected), statistics using ANOVA analysis with Bonferroni’s multiple comparison test (n=4 independent experiments)

**Supplementary Figure 4. Quantitative Mass spectrometry analysis of CRTC1 in EndoC-βH1 cells upon miR-184 inhibition + CRTC1 silencing. (a)** FASTA aminoacidic (aa) sequence of human CRTC1 protein; red highlighted text represents the peptide aa sequence analyzed in targeted Mass spectrometry (MS). (**b**) Representative raw spectrum of CRTC1 targeted MS in EndoC-βH1 cells transfected with synthetic inhibitor of hsa-miR-184-3p and CRTC1 siRNA. **(c)** Bar-dots plot graph showing quantization profile of CRTC1 expression by targeted MS in EndoC-βH1 after transfection with synthetic inhibitor of miR-184-3p or scramble inhibitor and with CRTC1 or CTR siRNA. Data are reported as mean±SD fold change vs control obtained by the ratio of NL values of CRTC1 normalized for NL values of both β-Actin and Chromogranin-A. Statistics performed using ANOVA analysis with Bonferroni’s multiple comparison test (n=4 independent experiments).

**Supplementary Figure 5. Representative cytofluorimetric plots and pyknotic nuclei count upon palmitate treatment and miR-184 inhibition + CRTC1 silencing in EndoC-βH1 cells. (a)** Representative cytofluorimetric plots of EndoC-βH1 cells treated or not with palmitate and transfected with synthetic inhibitor of miR-184-3p or scrambled miR-inhibitor and with CRTC1 or CTR siRNA**.** (**b**) Pyknotic nuclei count of EndoC-βH1 treated or not with palmitate and transfected with the synthetic inhibitor of miR-184-3p. Data are reported as mean±SD of the percentage of positive pyknotic nuclei cells on total nuclei. *p≤0.05 vs not treated, Scrambled miR-Inhibitor+Ctr siRNA; # p≤0.05 vs palm. treated, Scrambled miR-Inhibitor+Ctr siRNA); ┼ p≤0.05 vs palmitate treated, Scrambled miR-Inhibitor+CRTC1 siRNA), statistics using ANOVA analysis with Bonferroni’s multiple comparison test (n=3 independent experiments).

**Supplementary Figure 6. Representative cytofluorimetric plots and pyknotic nuclei count upon cytokines mix treatment and miR-184 inhibition + CRTC1 silencing in EndoC-βH1 cells. (a)** Representative cytofluorimetric plots of EndoC-βH1 cells treated or not with cytokines mix and transfected with synthetic inhibitor of miR-184-3p or scramble inhibitor and with CRTC1 or CTR siRNA**.** (**b**) Pyknotic nuclei count of EndoC-βH1 cells treated or not with cytokines mix and transfected with synthetic inhibitor of miR-184-3p. Data are reported as mean±SD of the percentage of positive pyknotic nuclei cells on total nuclei. *p≤0.05 vs not treated, Scrambled miR-Inhibitor+Ctr siRNA); #p≤0.05 vs cytokines treated, Scrambled miR-Inhibitor+Ctr siRNA. Statistics performed using ANOVA analysis with Bonferroni’s multiple comparison test (n=3 independent experiments).

**Supplementary Figure 7. NKX6.1 binding sites on human and murine miR-184 promoter sequence. (a)** Mouse miR-184-3p promoter sequence showed 1 predicted binding site for Nkx6.1 (green sequence, core matrix in bold); red sequence represents miR-184 gene sequence. **(b)** Human miR-184-3p promoter sequence showed 3 predicted binding site for NKX6.1 (green sequence, core matrix in bold); red sequence represents miR-184 sequence.

**Supplementary Figure 8. NKX6.1 nucleus-cytoplasm translocation reduces MafA mRNA expression in MIN-6 cells.** MafA mRNA expression evaluated through RT Real-Time PCR in MIN6 cells after treatment with 100 µM of H_2_O_2_ for 90. Data are reported as mean±SD of fold change. Statistics performed using paired Student’s t test (*p<0.05).

**Supplementary Figure 1**

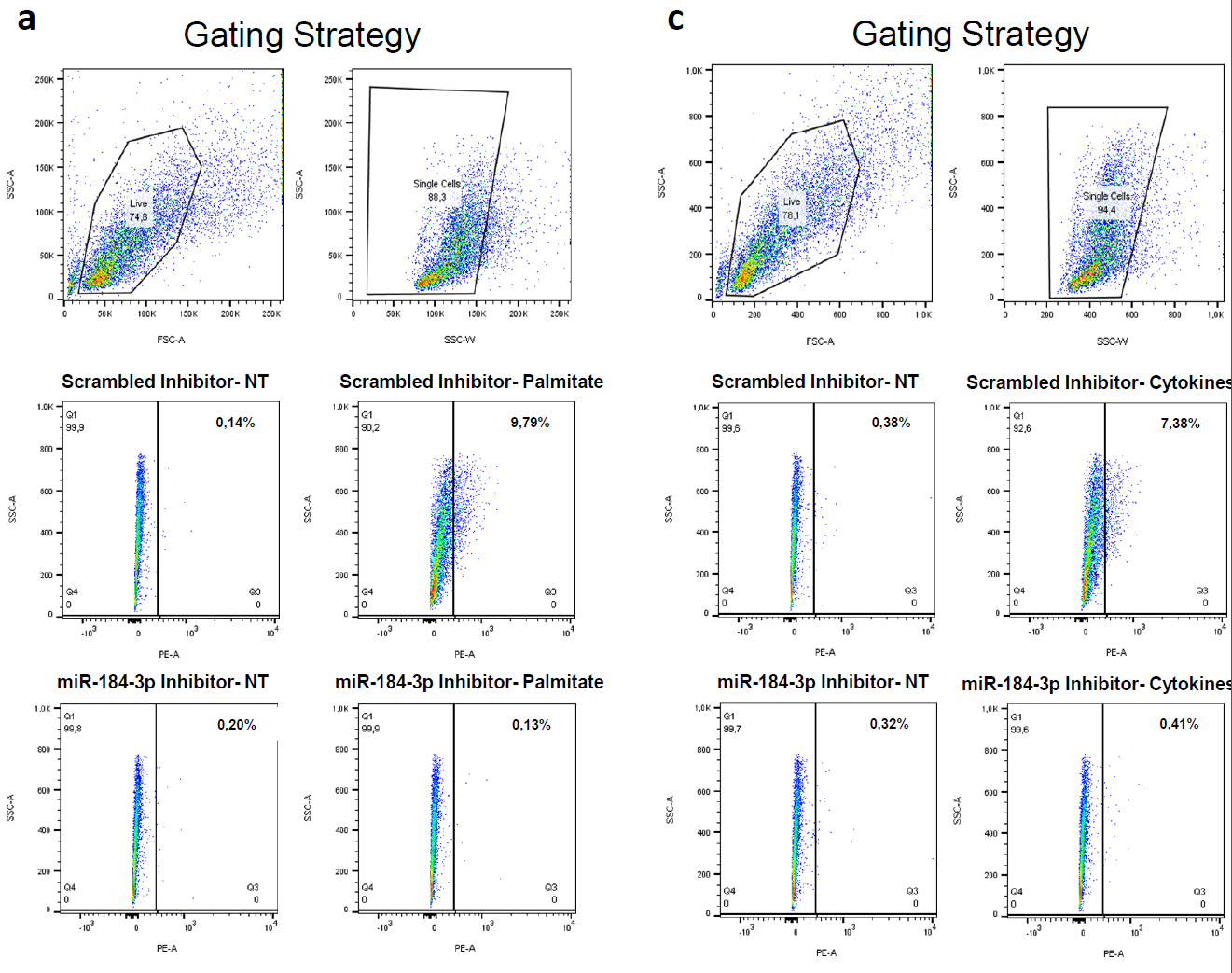

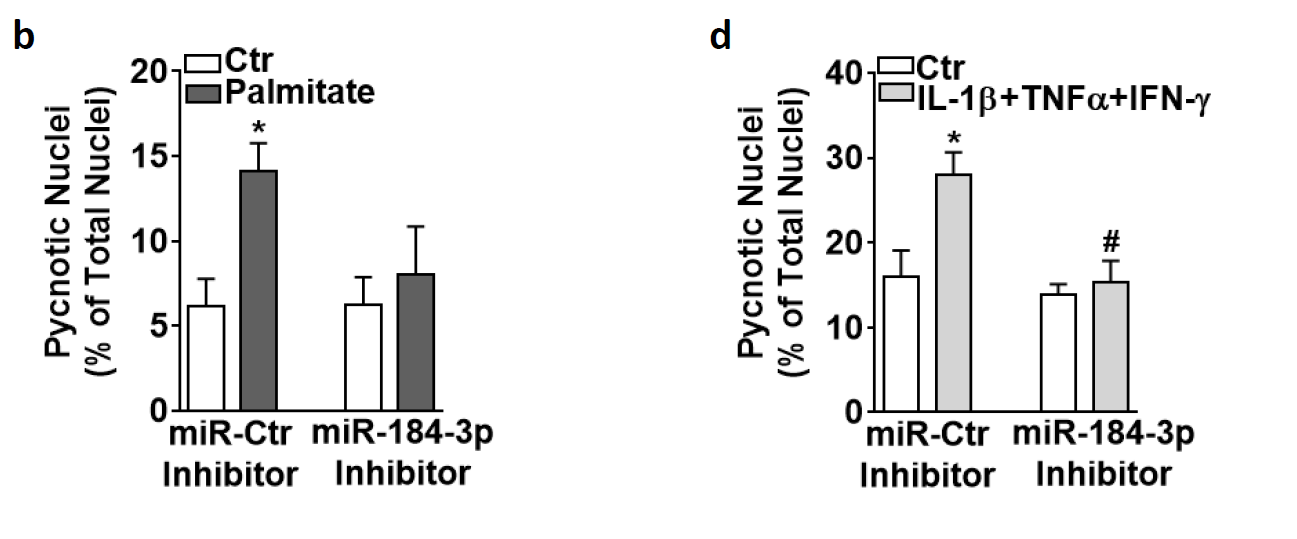

**Supplementary Figure 2**

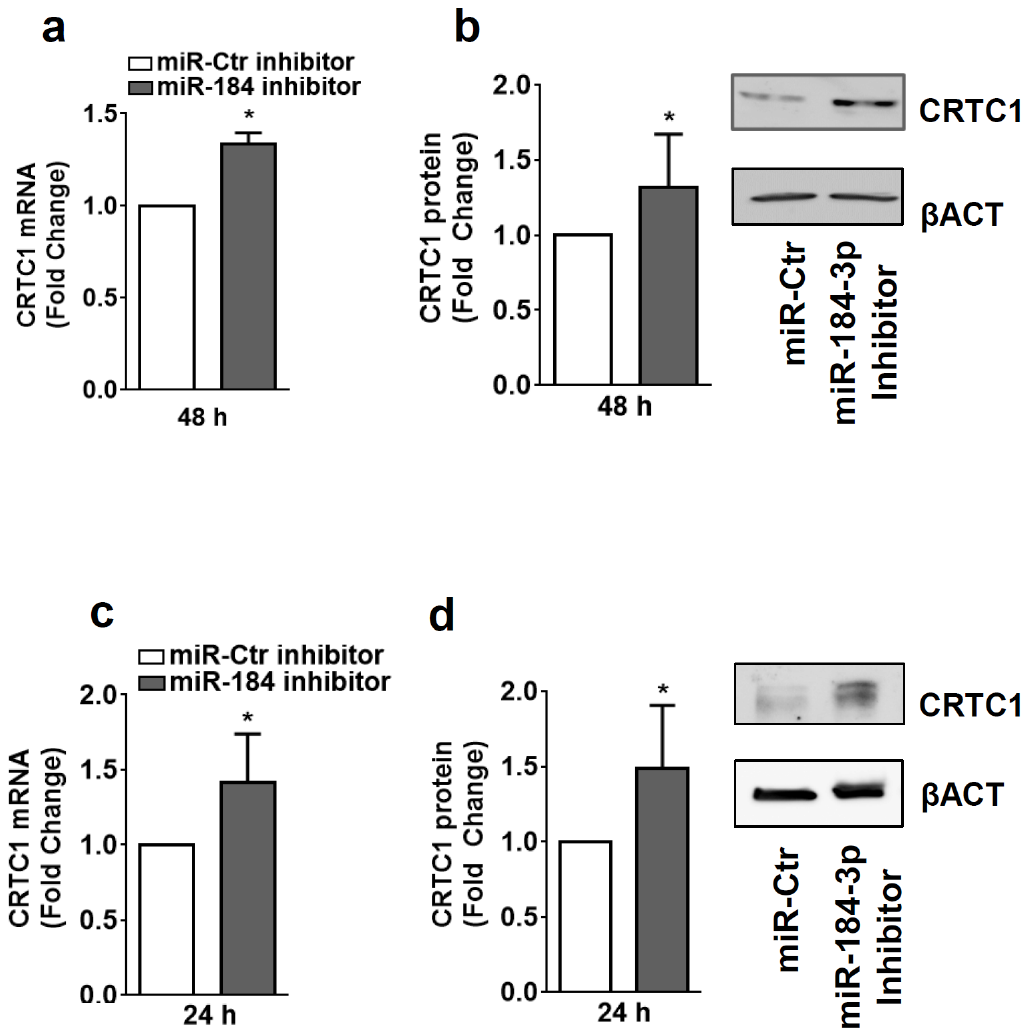

**Supplementary Figure 3**

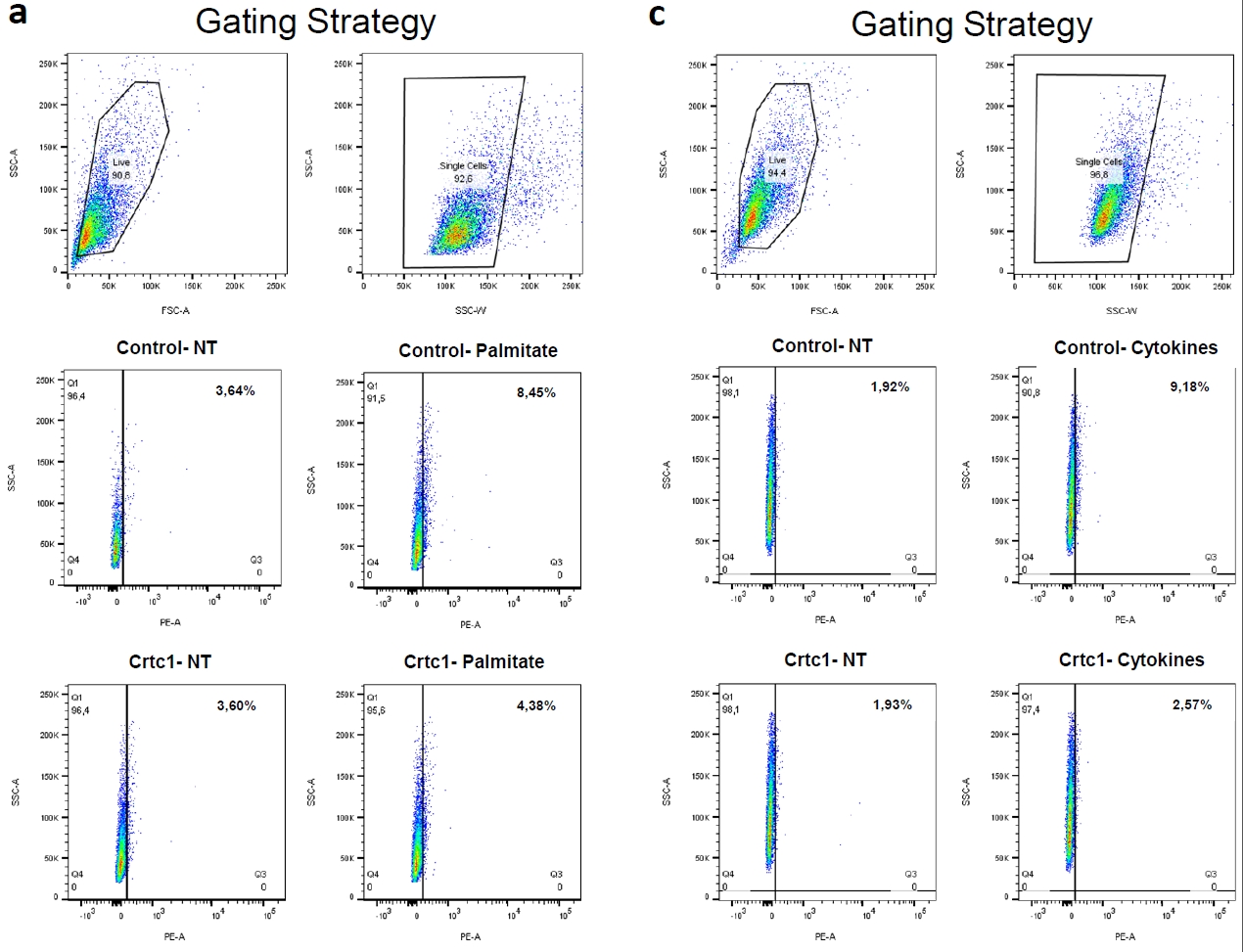

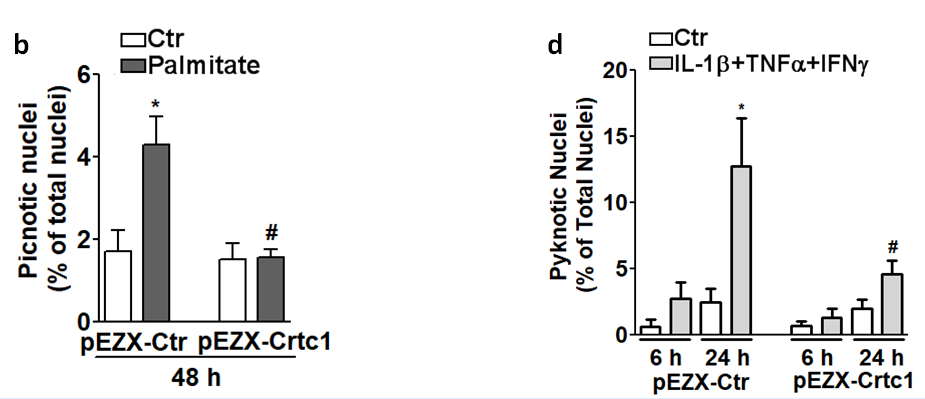

**Supplementary Figure 4**

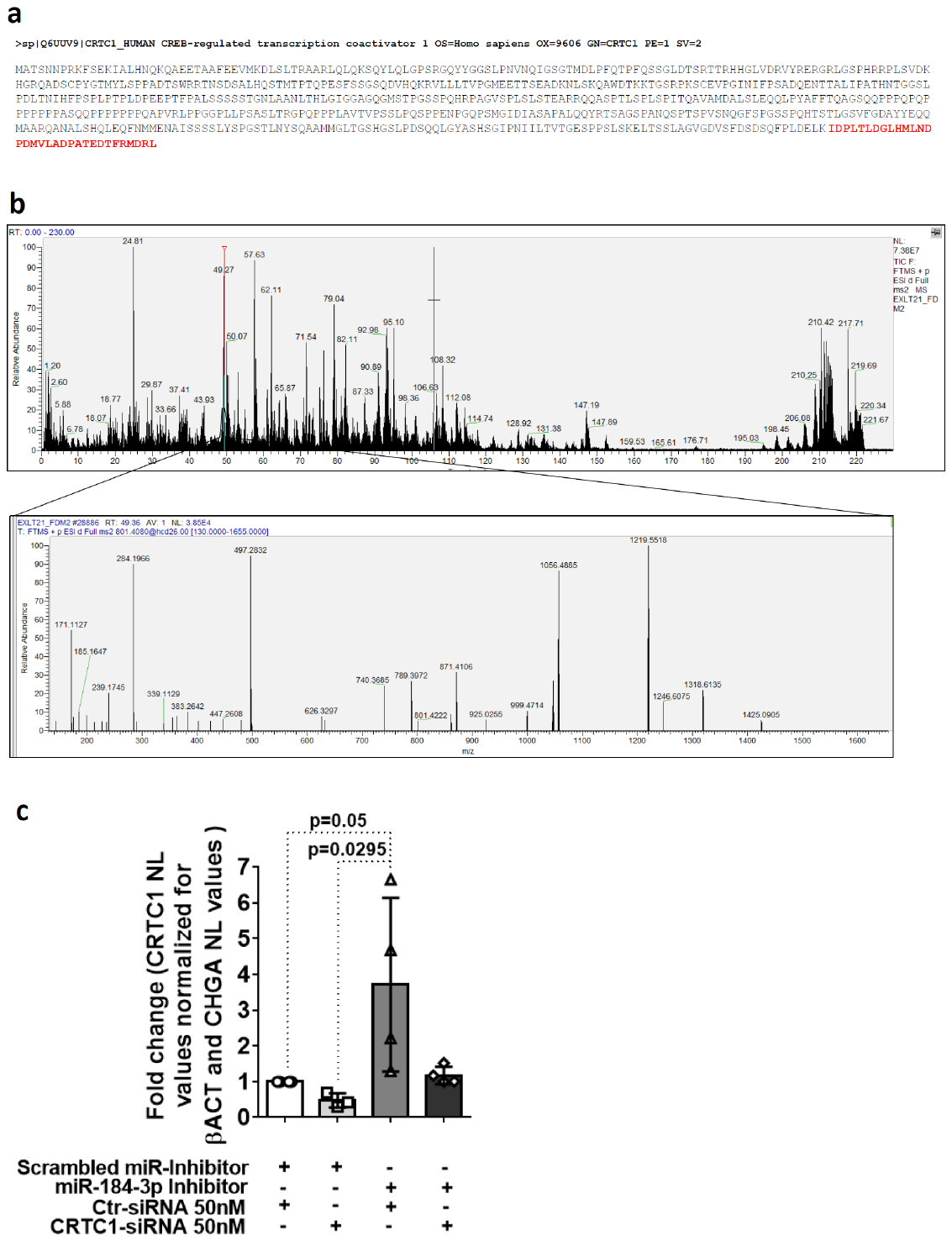

**Supplementary Figure 5**

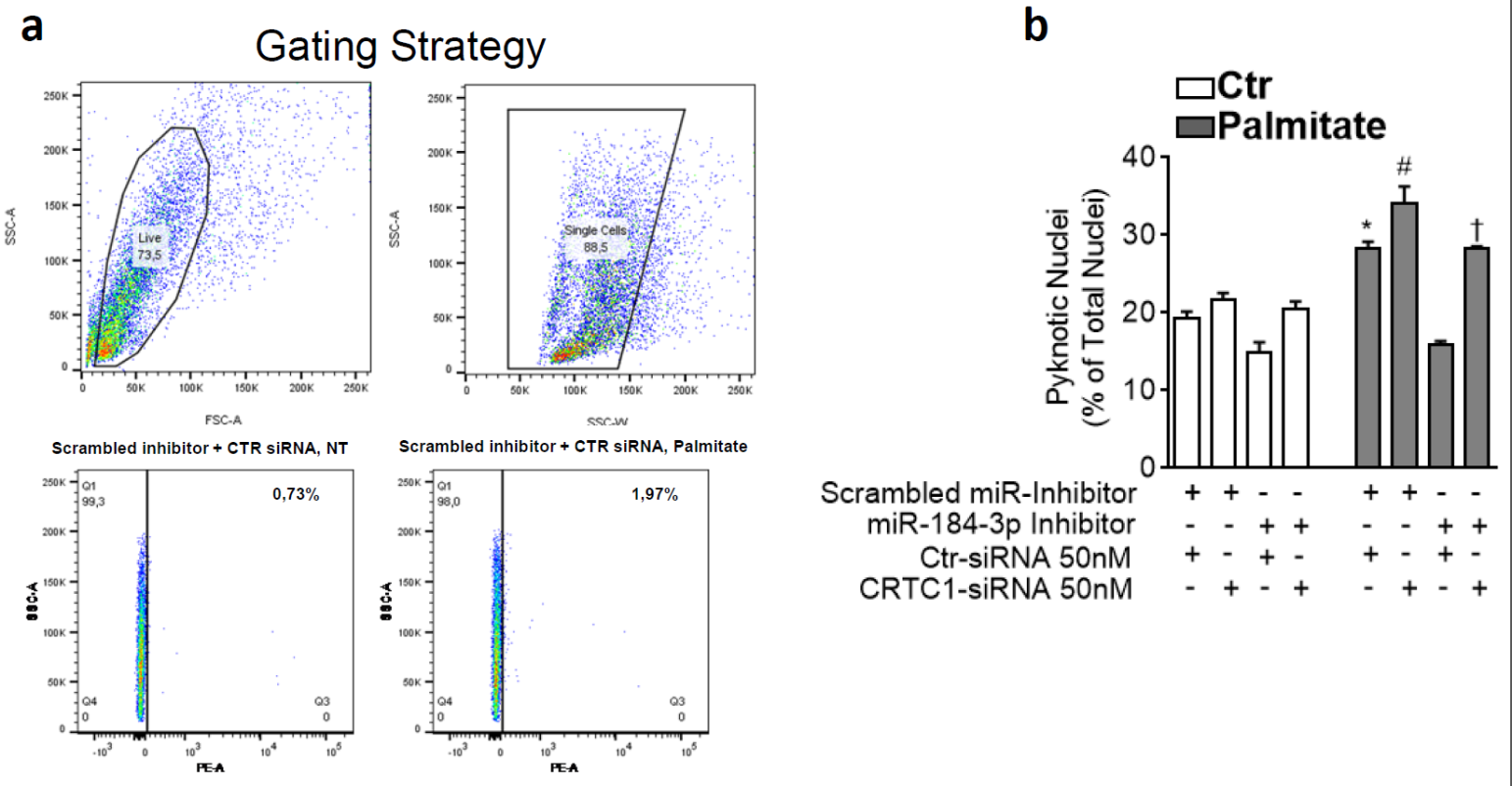

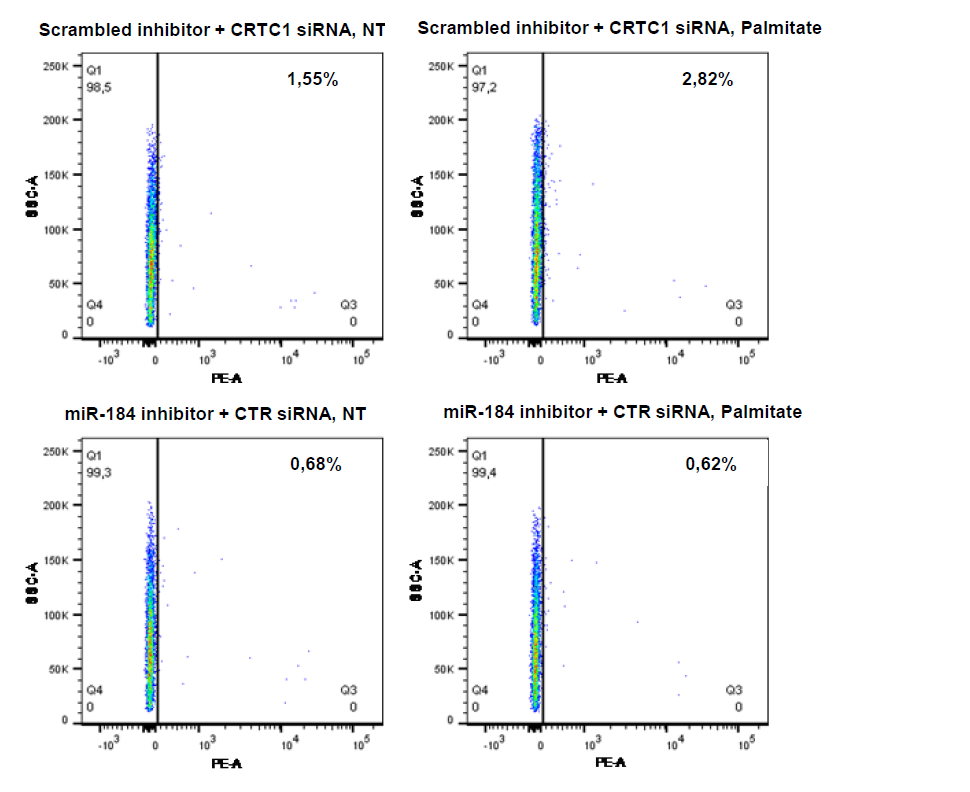

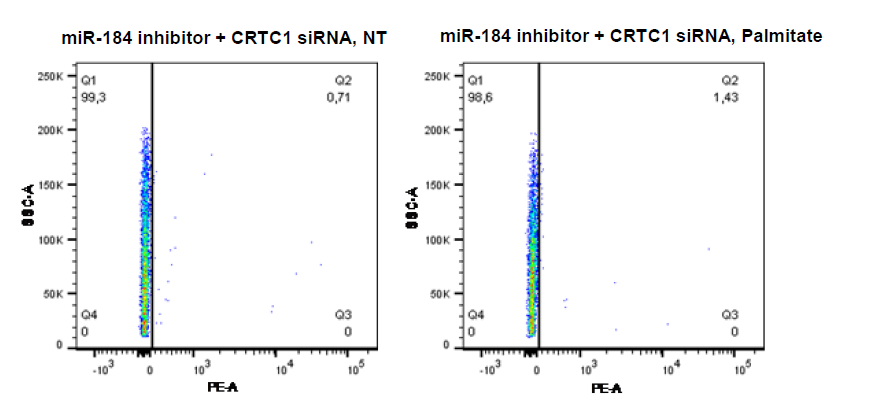

**Supplementary Figure 6**

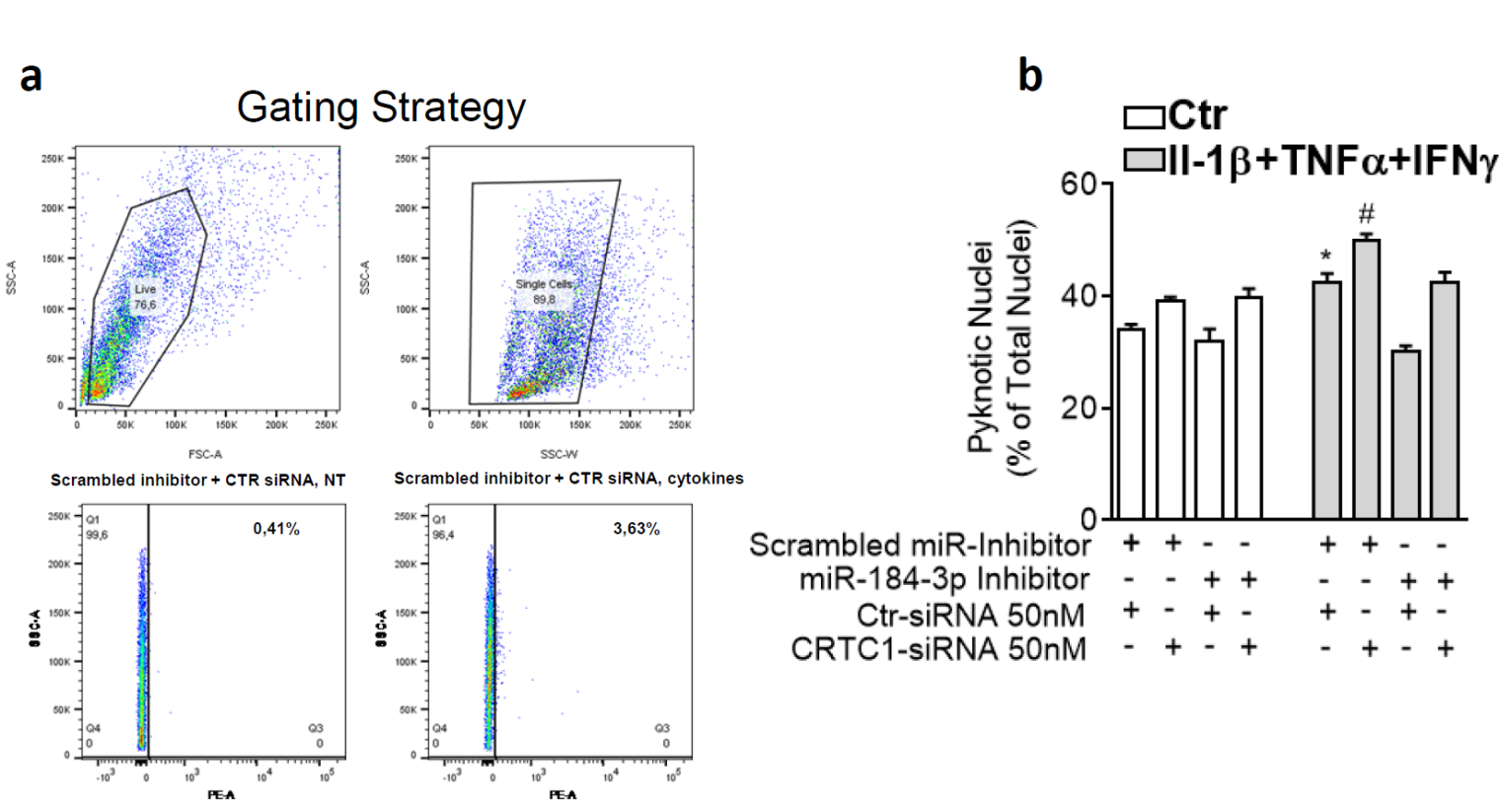

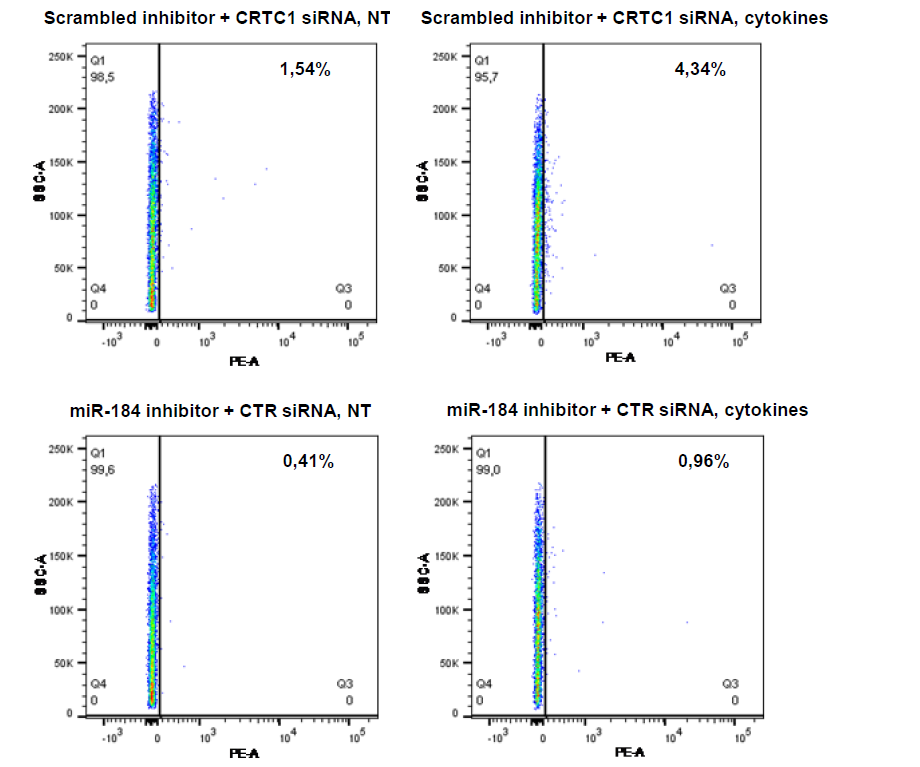

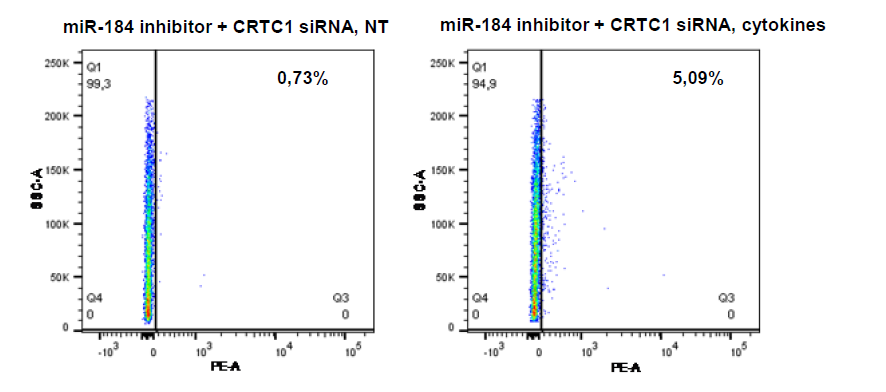

**Supplementary Figure 7**

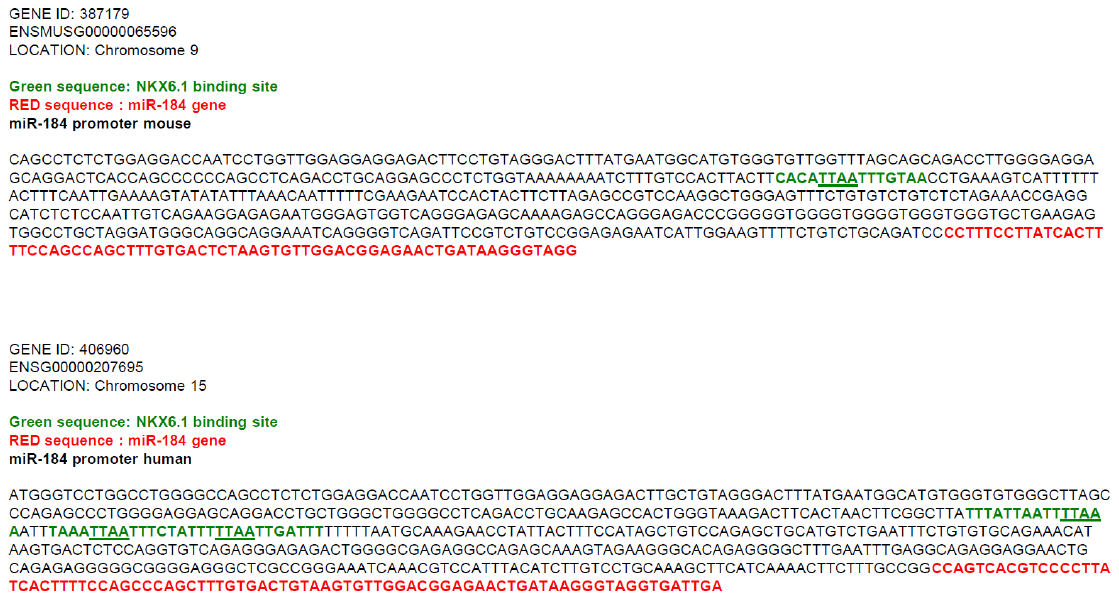

**Supplementary Figure 8**

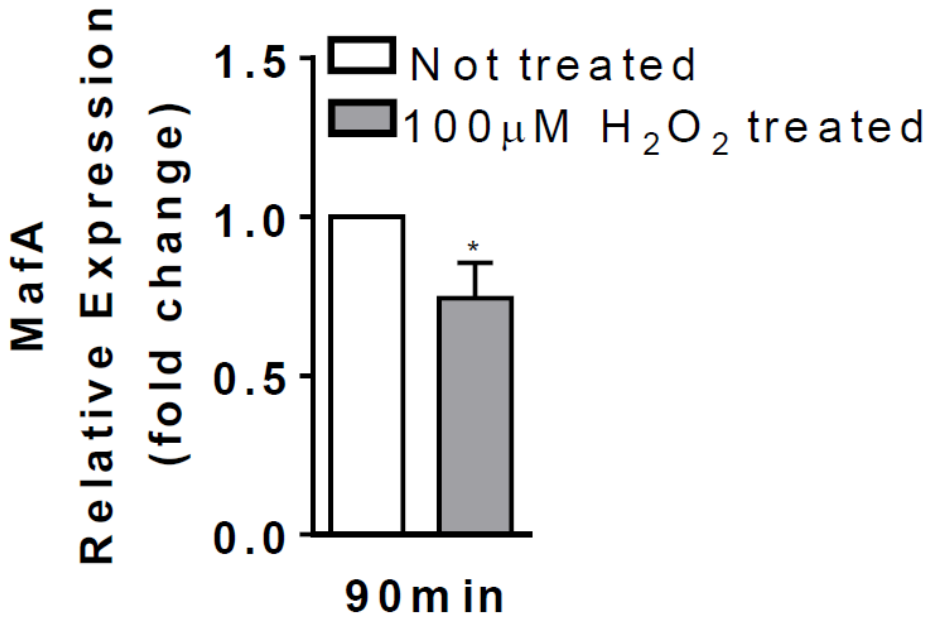
